# Nanoscale 3D DNA tracing reveals the mechanism of self-organization of mitotic chromosomes

**DOI:** 10.1101/2024.10.28.620625

**Authors:** Kai Sandvold Beckwith, Andreas Brunner, Natalia Rosalia Morero, Ralf Jungmann, Jan Ellenberg

**Affiliations:** Cell Biology and Biophysics, European Molecular Biology Laboratory, Heidelberg, Germany; Dept. Biomedical Laboratory Science, Norwegian University of Science and Technology, Trondheim, Norway; Collaboration for Joint PhD Degree between EMBL and Heidelberg University, Faculty of Biosciences, Heidelberg, Germany; Max Planck Institute of Biochemistry, Martinsried, Germany

## Abstract

How genomic DNA is folded during cell division to form the characteristic rod-shaped mitotic chromosomes essential for faithful genome inheritance is a long-standing open question in biology. Here, we use nanoscale DNA-tracing in single dividing cells to directly visualize how the 3D fold of genomic DNA changes during mitosis, at scales from single loops to entire chromosomes. Our structural analysis reveals a characteristic genome scaling minimum at 6-8 Mbp in mitosis. Combined with data-driven modeling and molecular perturbations, we can show that very large and strongly overlapping loops formed by Condensins are the fundamental structuring principle of mitotic chromosomes. These loops compact chromosomes locally and globally to the limit set by chromatin self-repulsion. The characteristic length, density and increasingly overlapping structure of mitotic loops we observe in 3D, fully explain how the rod-shaped mitotic chromosome structure emerges by self-organization during cell division.

## Introduction

Upon cell division, chromosomes undergo major shape changes into compact and stiff rod-like entities enabling their segregation into daughter cells^1-4^. Normal mitotic chromosome structure depends on loop-extruding Condensin complexes^5–8^. Studies using quantitative imaging of Condensins in single cells^9^ or biochemical cross-linking of genomic DNA followed by sequencing (HiC) in synchronized cell populations^10–12^ have hypothesized hierarchical looping models to underlie mitotic chromosome structure. According to these models, the two Condensin isoforms should fold chromosomal DNA initially into larger loops (∼500 kb) and then subdivide them into smaller (∼100 kb) loops. Early classical EM-studies on extracted chromosomes already suggested a loop-like organization of mitotic chromatin^13^, but the high density of DNA in mitotic chromosomes has precluded a direct visualization of its fold in natively fixed cells^14^. Therefore, the in situ internal organization of mitotic chromosomes remains unresolved, and it remains unclear how the higher-order functional properties of mitotic chromosomes for genome inheritance emerge from the molecular scale structure. Fluorescence based DNA tracing enables direct visualization of genome organization in single cells^15,16^ but has so far not been applied to mitotic chromosomes. We have previously established a mild, structure-preserving DNA tracing workflow (LoopTrace^17^), based on targeted single strand resection of DNA^18–21^, which allows high-fidelity nanoscale tracing of single chromatids in situ^17,22^. Here, we apply LoopTrace to visualize the organization of mitotic chromosomes from the kilobase to full chromosome scale in single dividing cells, and use this structural data for polymer simulations, allowing us to propose a self-organization model for mitotic chromosome folding.

## Results

### Multiscale DNA tracing from interphase to metaphase

To obtain the first direct view of chromosome refolding from interphase to metaphase, we designed FISH probe libraries for tracing entire human chromosomes (using chromosomes 2 and 14 as examples) at 1 Mb genomic resolution (Fig. 1A, B), 10 Mb regions at 200 kb resolution, as well as 1.2 Mb regions at 12 kb resolution. After refining our protocols for non-denaturing DNA tracing at the nanoscale^17^ for mitotic cells (Fig. S1A, B), we could routinely generate multiscale chromosome traces from interphase to metaphase (Fig. 1C). Importantly, sequential imaging of the 12 kb resolved regions achieved a median of 85% trace-completeness and better than 25 nm 3D precision, sufficient to reliably resolve individual DNA loops (estimated to be ∼100-500 kb^9,11^, Fig. S1C, D). As expected, interphase chromosomes appeared overall as distinct ellipsoid territories within the cell nucleus^23^ and DNA looping structures consistent with known TADs were readily visualized at 12 kb resolution (Fig. 1C, Fig. S1E, F). Upon entry into mitosis such genomically reproducibly positioned clusters of small loops were no longer present and instead variably positioned, longer and increasingly backfolded DNA loops appeared (Fig. 1C prophase to metaphase at 1.2 Mb and 10 Mb resolutions). At the same time, chromosomes started to individualize into long, thin and often irregularly bent and variably compacted structures (Fig. 1C). During prometaphase, chromosomes became more contiguously axially compacted, when their arms thickened and straightened, which continued until they reached a compact, rod-like and often overall V-like shape in metaphase (Fig. 1C, Fig. S1G, H).

**Figure 1.**
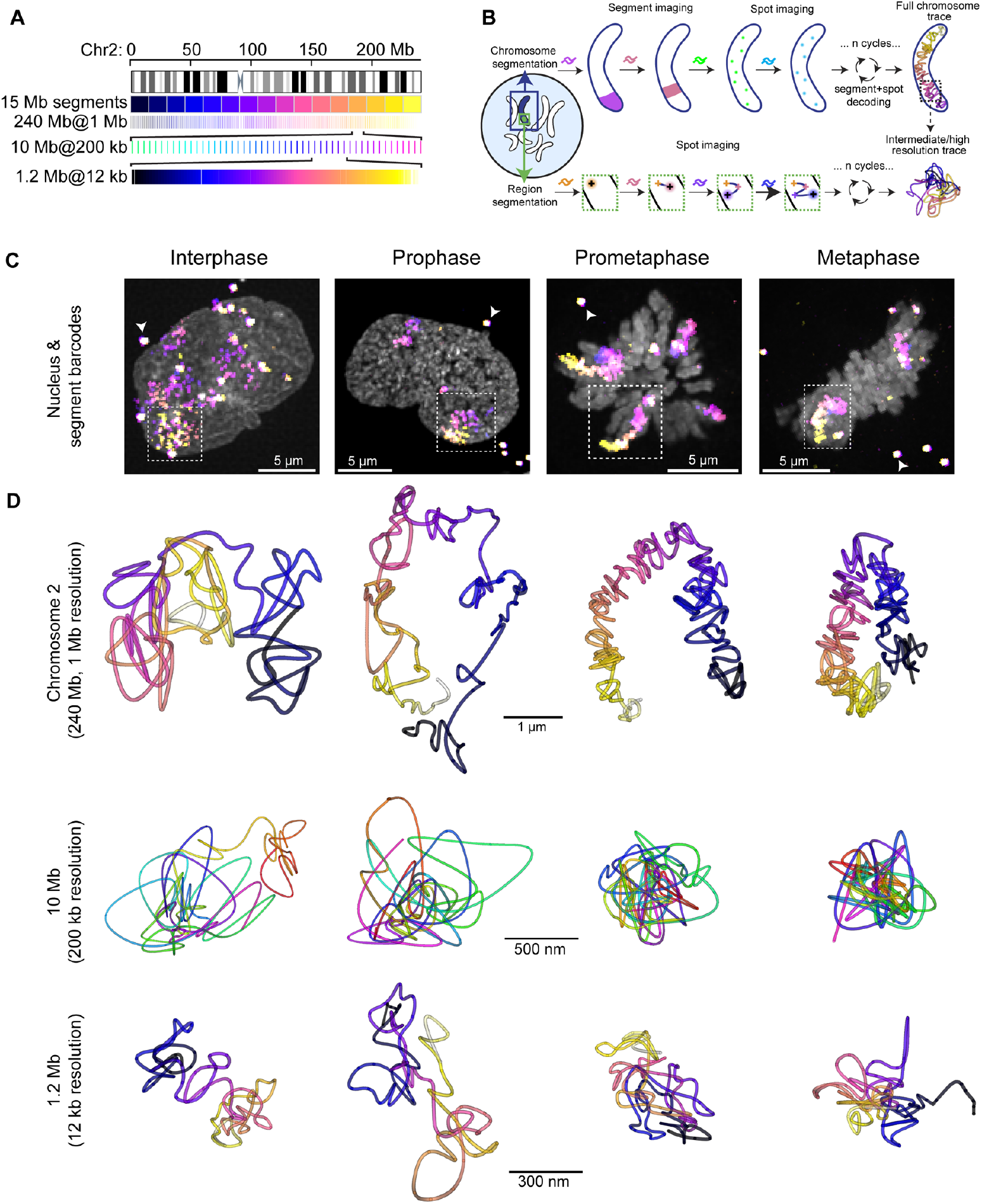
Multiscale DNA tracing from interphase to metaphase. **A** Schematic of multiscale probe libraries targeting chromosome 2, which was decoded using a combination of 15 Mb segment barcode and 1 Mb resolution spot barcodes. Full chromosome libraries were supplemented with intermediate scale (30 kb spots every 200 kb for 10-12 Mb) and high resolution (12 kb spots contiguously tiled for 1.2 Mb) libraries targeting the same and different (Chr5, Chr14, Chr18) chromosomes. **B** Schematic of multiscale chromosome trace acquisition. Whole chromosomes were segmented based on the full library signal, while sequential images of 15 Mb segments and 100 kb spots were used to decode and uniquely identify and 3D localize all detected spots. Intermediate and high-resolution libraries were segmented by regional barcodes and traced by sequential spot fitting in 3D. **C** Exemplary maximum-intensity projected micrographs of cell nuclei (DAPI, grey) with full chromosome traces (color-coded sequential 15 Mb segments). Arrowheads indicate fiducial beads used for drift correction. **D** Reconstructed 3D traces (multi-coloured) from libraries targeting whole chromosome 2 and intermediate- and high-resolution regions. Data representative of in total 8879 traces in 1396 cells in 5 independent experiments.

Having 3D traces from hundreds of mitotically staged single cells allowed us to quantitatively analyze changes in the DNA fold during mitosis, including the abundance, size and internal nesting of loops, as well as the radius of gyration as a general measure of chromosome compaction (Fig. 2A, Fig. S2A). Interestingly, mitotic chromosomes exhibited a significant increase in the number and size of loops from prophase onwards (Fig. 2B, Fig. S2B-D). This was followed by an increase in nesting of loops with overlapping bases (Fig 2B, Fig. S2A loop measurement illustration, Fig. S2B-D) starting in prometaphase when the major wave of Condensin I binding occurs^6,9,24^. Nesting expectedly coincided with a significant decrease of the radius of gyration, indicating that loop-scale compaction occurs mainly in prometaphase (Fig. 2B, Fig. S2B-D). However, the decrease of the radius of gyration began already in prophase, indicating that both loop size increase and loop backfolding and nesting underlie mitotic chromosome compaction. Analyzing these parameters revealed that the structural features that change during mitosis are well sampled at this genomic resolution (Fig. S2B-D).

**Figure 2.**
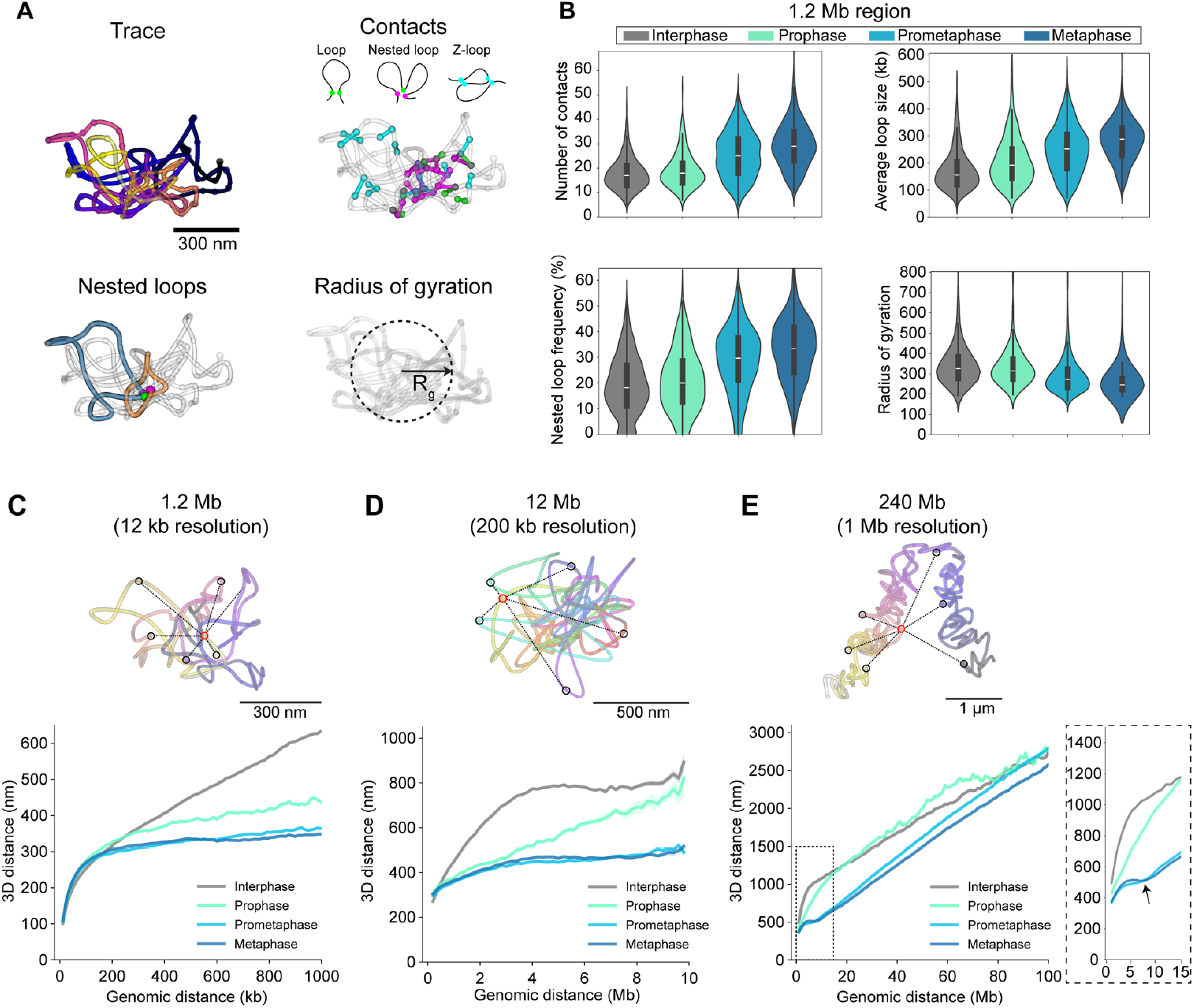
Mitosis-specific chromatin loop and distance scaling signatures. **A** Illustration of single-trace metrics. Euclidean distance-based filtering of close contacts (<100 nm) with a genomic distance over 30 kb defines “loops”. Based on their overlap with other loops or loop bases, contacts are classified into base loops, nested loops (when 2 or more base loops coincide to form a second, larger loop) or z-loops (2 loops partially overlap). Radius of gyration is a measure of overall compaction of the trace. **B** Trace metrics calculated for the high-resolution (12 kb) 1.2 Mb region on chromosome 5 (149,500,723-150,699,962). Trace metrics were calculated from traces that were more than 80% complete. Data from n=278 (510), n=22 (44), n=124 (297) and n=152 (398) inter-, pro-, prometa- and metaphase cells (traces) from 3 independent experiments. Median, quartiles and whiskers are shown in the plots. **C, D, E** Traces of a high-resolution region (chromosome 5, C), intermediate scale (chromosome 2, B) and whole chromosome 2 (E) and corresponding pairwise distance scaling plots. Exemplary pairwise distances (black) for a single spot (red) within the DNA trace are highlighted. Inset with arrow highlights the characteristic scaling dip at 6-8 Mb. Plots show median ± standard error of the mean. Data from (C) n=400 (824), n=39 (109), n=241 (609) and n=244 (644), 3 experiments; (D) n=165 (237), n=27 (44), n=175 (325) and n=187 (346); 2 experiments; and (E) n=331 (803), n=28 (83), n=241 (736) and n=212 (686), 3 experiments; inter-, pro-, prometa- and metaphase cells (traces).

The multiscale DNA tracing of mitotic chromosomes also provided an opportunity to examine the question of higher order regularity of the DNA fold ^2^. We did not observe a highly regular organization at either the whole chromosome, the 10 Mb regions or at the scale of single loops within 1 Mb subregions, suggesting that loops are stochastically organized relative to the geometric chromosome axis (Fig. 1C, metaphase). Consistently, statistical analysis of symmetry parameters relative to the central axis did not show significant helicity at the full chromosome scale (Fig. S2E).

### Mitotic chromosome scaling dips at 6-8 Mb

If mitotic chromosomes become more compact at each scale, their genomic versus physical 3D distance scaling should also change from interphase to mitosis. To probe distance scaling of chromosomal DNA for additional structural features that might arise during mitosis, we plotted all pairwise Euclidian distances versus the respective pairwise genomic distance for all traces obtained at the three genomic scales, grouped by cell cycle stage (Fig. 2C-E). This analysis revealed a characteristic scaling behavior of mitotic chromosomes. As expected from the increase in close contacts, mitotic chromosomes showed a significantly higher compaction at the 200 kb - 1Mb scale starting in prophase, which was essentially completed in prometaphase (Fig. 2C). Interestingly, this correlated with a slight decompaction of chromatin at short genomic distances of less than 200 kb (Fig. 2C, Fig. S2F, G). At the 1 - 10 megabase scale, mitotic chromosomes showed an increasing long-range compaction in prophase to prometaphase (Fig. 2D, Fig. S2F, G).

Not predictable from the loop-scale physical distance metrics, the scaling analysis revealed a higher order local minimum at 6-8 megabases indicating increased backfolding of chromatin. This scaling “dip” at large genomic distance appeared in prometaphase and became even more pronounced in metaphase (Fig. 2D, Fig. S2F, G). It was also clearly present at the whole chromosome scale (Fig. 2E, Fig. S2F), showing that it is a general structural feature and not specific to the three analyzed 10 Mb regions. The whole-chromosome scaling plot furthermore showed that while chromosomes are already significantly more compact in prophase than in interphase at a scale below 10 Mb, they are not yet maximally compacted along their entire length and display a rather irregular scaling behavior above 10 Mb early in mitosis (Fig. 2E). By contrast, in prometa- and metaphase, global chromosome compaction is largely completed, with a smooth linear scaling behavior above the 6-8 Mb mitotic dip.

### Mitotic chromosome structural features depend on Condensins

The presence of the directly traced, on average 300 kb long, mitosis-specific loops that became increasingly nested would be expected to be the result of loop extrusion activity by Condensin complexes^9,11^. To test which structural features of mitotic chromosomes depend on loop extrusion across the sampled scales, we depleted Condensins prior to mitosis in HeLa Kyoto cells with the Condensin subunit SMC4 endogenously tagged with an auxin-inducible degron in all alleles ^25^. We then performed multiscale chromatin tracing of Condensin-depleted metaphase cells (Fig. S3A) and analysed the 3D trace parameters as well as their genomic-physical distance scaling (Fig. 3A-D, Fig. S3B, C). As expected, Condensin depletion resulted in a dramatic loss of individual mitotic loops and their nesting and a corresponding decrease in number of close contacts, demonstrating that mitotic DNA loops and their nesting depend on Condensin (Fig. 3B). Strikingly, when we examined the distance scaling relationship, the characteristic mitotic dip at 6-8 Mb was also completely abolished in ΔSMC4 chromosomes (Fig. 3D, Fig. S3C). In addition, the smooth linear distance scaling behavior above 10 Mb became rather variable after Condensin depletion, consistent with the irregular and no longer clearly rod-like shape chromosomes exhibited in the corresponding 3D traces of entire chromosomes (Fig. 3C). Consistently, ΔSMC4 chromosomes also failed to reduce their radius of gyration as a measure of general compaction (Fig. 3B). Thus, not only individual mitotic loops and their internal nesting, leading to local compaction, but also the characteristic mitotic 6-8 Mb dip and the compaction of entire chromosomes depend on Condensins.

**Figure 3.**
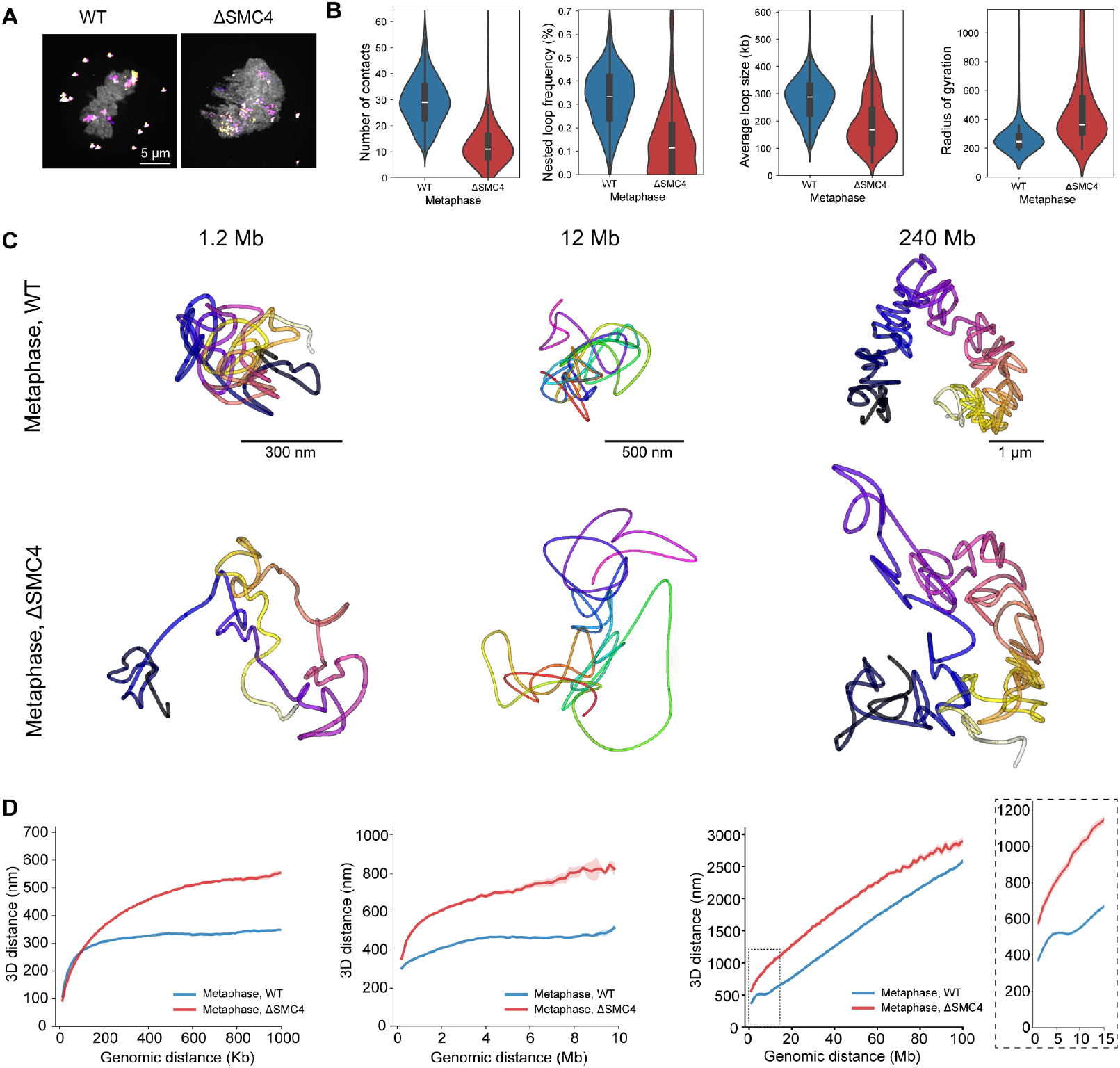
Mitotic chromosome structural features depend on Condensins. **A** Wild type (WT) HeLa Kyto cells and HeLa Kyoto cells with SMC4-mAID-Halo acutely depleted for 3-4 hours with 5-Ph-IAA before mitotic entry (ΔSMC4). Exemplary maximum-intensity projected micrograph of metaphase chromatin (DAPI, grey) with full chromosome 2 traces (DNA-FISH, multi-coloured). Data representative of 212 WT metaphase traces from 3 independent experiments and 84 metaphase ΔSMC4 traces from 2 independent experiments. **B** Trace metrics from chr5:149500723-150699962 (1.2 Mb, 12 kb resolution) for WT and ΔSMC4 cells. Data from 152 (398) WT cells (traces), 3 independent experiments, and 153 (265) ΔSMC4 cells (traces), 2 independent experiments. **C** 3D DNA traces for chromosome 5, 1 Mb scale (12 kb resolution), chromosome 2, 10 Mb scale (200 kb resolution) and whole chromosome 2 (1 Mb resolution) from WT and ΔSMC4 cells at metaphase. Data representative of Chr5, 1 Mb: 152 (398) WT cells (traces), 3 independent experiments, and 153 (265) ΔSMC4 cells, 2 independent experiment; Chr 2, 10 Mb: 159 (286) WT cells (traces), 2 independent experiments and 106 (153) ΔSMC4 cells (traces), 2 independent experiments; Chr2 whole: 130 (212) WT cells (traces), 3 independent experiments and 51 (62) ΔSMC4 cells (traces), one experiment. **D** Distance scaling plots for chromosome 5, 1 Mb scale, chromosome 2, 10 Mb scale and whole chromosome 2 at metaphase. Data from Chr5, 1 Mb: 244 (644) WT cells (traces), 3 independent experiments and 212 (608) ΔSMC4 cells (traces), 2 independent experiments; Chr2, 10 Mb: 187 (346) WT cells (traces), 3 independent experiments and 124 (193) ΔSMC4 cells (traces), 2 independent experiments; Chr2, whole: 212 (686) WT cells (traces), 3 independent experiments and 206 (503) ΔSMC4 cells (traces), 2 independent experiments.

### Loop extrusion and self-repulsion correctly predict mitotic chromosome structure

How then could micrometer-scale rod-shaped mitotic chromosomes arise from molecular scale architecture of individual DNA loops? Previous theoretical work has shown that, in principle, a self-avoiding polymer fibre constrained by cross-links^26^ or loops^27^ could form a rod-like shape through entropic self-repulsion. This is because crosslinks/loops backfold the fibre on the one hand, while on the other hand the increased density of loops leads to more frequent collisions and a self-repulsion of loops that seeks to maximize the polymer’s available conformational space^26–29^.

While it is clear from our data (Fig. 3) and previous work^4^ that the Condensin complexes are required for the formation of stiff mitotic chromosome rods, if the generation of loops alone is sufficient to explain our newly determined 3D structure and scaling properties of mitotic chromosomes remained unclear. To test this mechanistically, we set up a polymer model of mitotic chromosomes constrained by the currently available quantitative experimental data on Condensins (see below) to compare it to our DNA tracing data. Building on pioneering work by the Mirny group^30^ and their open-source simulation framework (https://github.com/open2c/polychrom), we simulated a realistic human chromosome of 100 Mb as a chain of 100,000 beads (e.g. 1 kb/bead), ensured volume exclusion through repulsion between monomers. and physically scaled the unconstrained polymer by high resolution ΔSMC4 traces (Fig S3A). We dynamically simulated Condensin-driven loop extrusion by the sequential action of initially two stably bound Condensin II complexes per megabase for 10 minutes (prophase to nuclear envelope break down) and then ten dynamically bound Condensin I complexes for an additional 30 minutes (prometa-to metaphase)^9,31^ (Supplementary Table S1^22^). Where available, we used parameters for Condensin abundance and DNA residence time determined in vivo, and used single molecule in vitro measurements for their uni-directional extrusion rate and high passing frequency^8,32^. Otherwise, our model was assumption-free especially regarding the structure of the polymer.

We found that this realistic and data driven polymer model has an excellent fit to our tracing data across the three experimentally measured scales from prophase to metaphase with an extrusion rate of 4±2 kbps and free passing of Condensins (Fig. 4A-D; Fig. S4B-E; Supplementary Movie S1). The simulation shows that Condensin-driven loop extrusion is sufficient to generate the experimentally observed mitotic chromosome fold and overall shape. Importantly, the simulation shows that the number of Condensin II complexes present in these cells, that are stably bound throughout mitosis, can extrude overlapping loops of on average 6-8 megabases when they freely pass each other. The size and overlapping nature of these loops directly explains the distance scaling minimum we observed in pro-/metaphase. By contrast, the number of Condensin I complexes present in cells, that are bound only for minutes, lead to nesting inside these large loops, thereby increasing the DNA density and number of loops and thus their self-repulsion. Our simulations furthermore show that loop extrusion and chromatin self-repulsion are sufficient to induce a straightening of the mitotic chromosome rod as its length shortens and its width increases (Fig. 4A), which is caused by increased self-repulsion of DNA at the concave side of the bent chromosome. Thus, the simulation explains the rod-shape and rigidity of chromosomes by a balance of the long-distance compaction forces of large, overlapping loops and the self-repulsion of these loops due to the entropic resistance of the polymer chain to be packed too densely.

**Figure 4.**
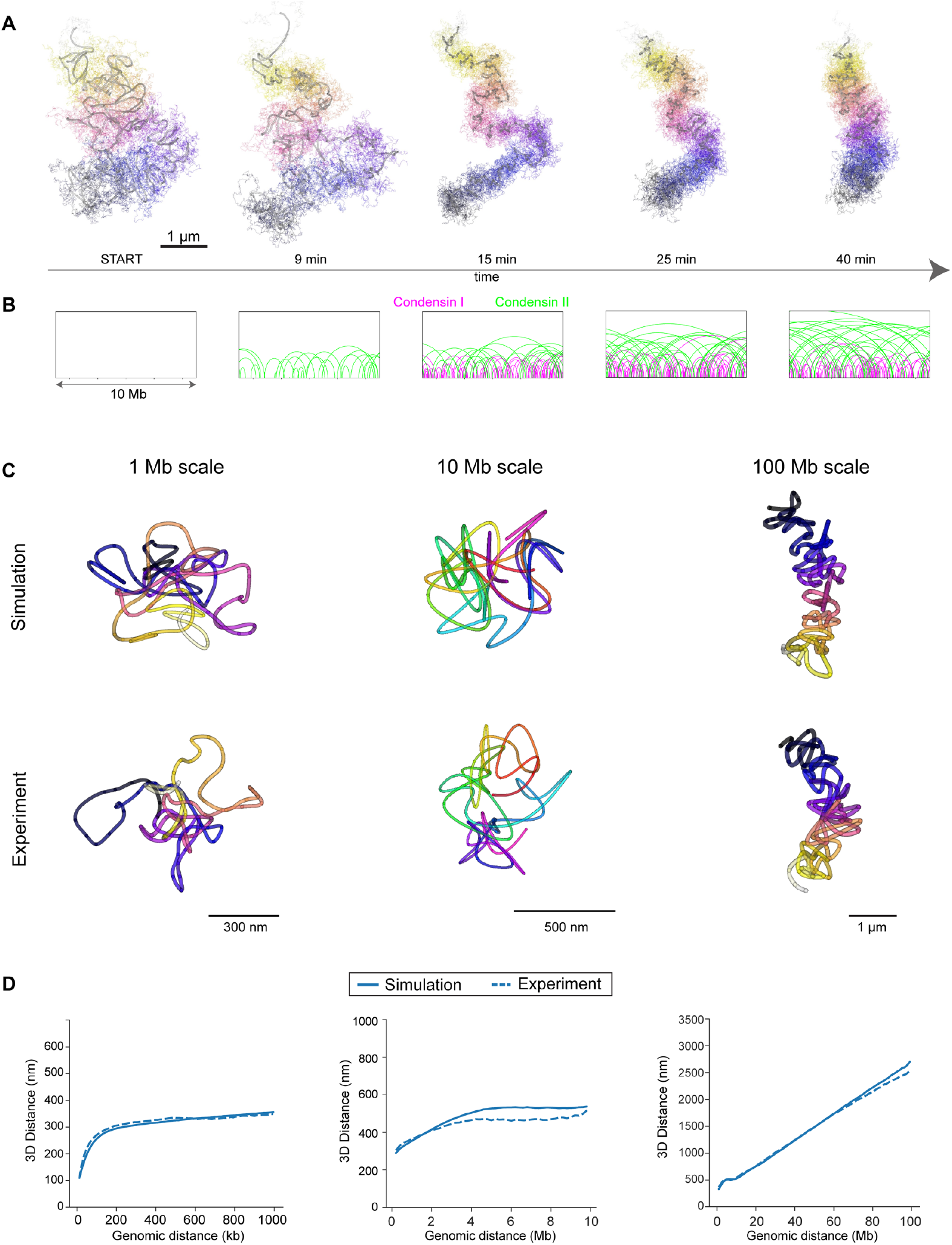
Loop extrusion and self-repulsion correctly predict mitotic chromosome structure. **A** Polymer simulation of mitotic progression for a 100 Mb chromosome sampled at the indicated timepoints, displayed at 2 kb resolution (multi-colored, thin line) and as a 1 Mb rolling average (grey, thick line). Data representative of 20 dynamically simulated chromosomes. **B** Condensin I (magenta) and Condensin II (green) loop lengths shown for a 10 Mb stretch of the simulation example shown in **A**. Loop height scales with length for clarity. **C** Reconstructed traces from simulated regions from 100 Mb chromosomes at metaphase (40 minutes) and corresponding experimental data (Chr5 1 Mb scale, Chr2 10 Mb scale, Chr2 100 Mb from q-arm). Simulated traces were sampled as experimental data (12 kb tiled probes, 30 kb probes with 200 kb resolution and 100 kb probes with 1 Mb resolution). Simulated data representative of 20 simulated 100 Mb chromosomes, experimental data representative of Chr 5, 1 Mb: 152 (398) cells (traces); Chr 2, 10 Mb: 159 (286) cells (traces); Chr2, 100 Mb: 130 (212), 3 independent experiments. **D** Distance scaling plots for simulated metaphase chromosomes and corresponding experimental data. Simulated chromosomes were sampled as in **C** either 10 times (10 and 100 Mb scales) or 60 times (1 Mb scale) with different starting positions in each of twenty 100 Mb simulated chromosomes for representative sampling. Experimental data from Chr5, 1 Mb: n=244 (644) cell (traces), 3 independent experiments; Chr2, 10 Mb: n=187 (346) cell (traces), 2 independent experiments; Chr2q, 100 Mb: n=212 (686) cell (traces), 3 independent experiments.

### The extrusion-repulsion model predicts realistic Condensin localization

If the core mechanism of self-organization is indeed Condensin driven loop extrusion, our polymer model should also be able to predict other structural features of mitotic chromosome architecture. We asked the simulation to output the predicted sub-chromosomal localizations of the two types of Condensin complexes in metaphase chromosomes. Without prior assumptions on Condensin positioning, the model predicted that Condensin II should localize more centrally with the chromosome, while Condensin I should localize more peripherally (Fig. S4F, G). Strikingly, these predictions match previous experimental measurements of Condensin isoform localizations within the chromosome by super-resolution microscopy^9^.

We can conclude that self-organization of chromosomal DNA by extrusion of long overlapping loops by Condensin II, nested by abundant and dynamic short-range loops by Condensin I and their self-repulsion is sufficient to explain how the experimentally observed structural and higher order emergent properties of mitotic chromosomes are generated.

### Global compaction is influenced by self-repulsion

In addition to loop extrusion, chromatin modifications have been reported to influence global mitotic chromosome organization^25^. We therefore asked if we could accurately predict the effect of a change in chromatin modifications, independent of changes to loop extrusion. We thus modeled an experimentally accessible chromatin modification, i.e. the net negative charge of histones, which can be changed with inhibition of histone deacetylases (TSA), which leads to hyperacetylation of normally positively charged lysines on histone tails^33^. More negatively charged chromatin should result in a higher self-avoidance, due to electrostatic and/or steric effects^34,35^. To predict the effect of higher self-avoidance in our model, we increased the monomer repulsion in our simulated polymer and observed a uniform expansion of mitotic chromosomes (Fig. 5A-E). Importantly, the model predicted that this global expansion of chromatin can occur without affecting the Condensin driven loop organization, and thus predicts that the main structural effect of higher self-repulsion should be to limit the density to which DNA loops can be packed. To validate this prediction experimentally, we performed mitotic chromosome tracing in cells in which histone deacetylase had been inhibited with TSA. TSA treated cells indeed exhibited unchanged Condensin-driven structural features such as loop abundance, size and nesting, as well as a preserved characteristic scaling dip at 6-8 Mb (Fig. 5A-E, Fig. S5A-D). By contrast, TSA-treated mitotic chromosomes exhibited a clear, multiscale decompaction of chromatin (Fig. 5D, Fig. S5C), leading to a 20% expansion of chromatid width and length, consistent with model predictions (Fig. 5E, Fig. S5D, E). Thus, our model can discriminate the distinct contributions of loop extrusion and chromatin state in shaping mitotic chromosomes, providing a mechanistic explanation for the observation that TSA treatment leads to global decompaction of mitotic chromosomes ^25^.

**Figure 5.**
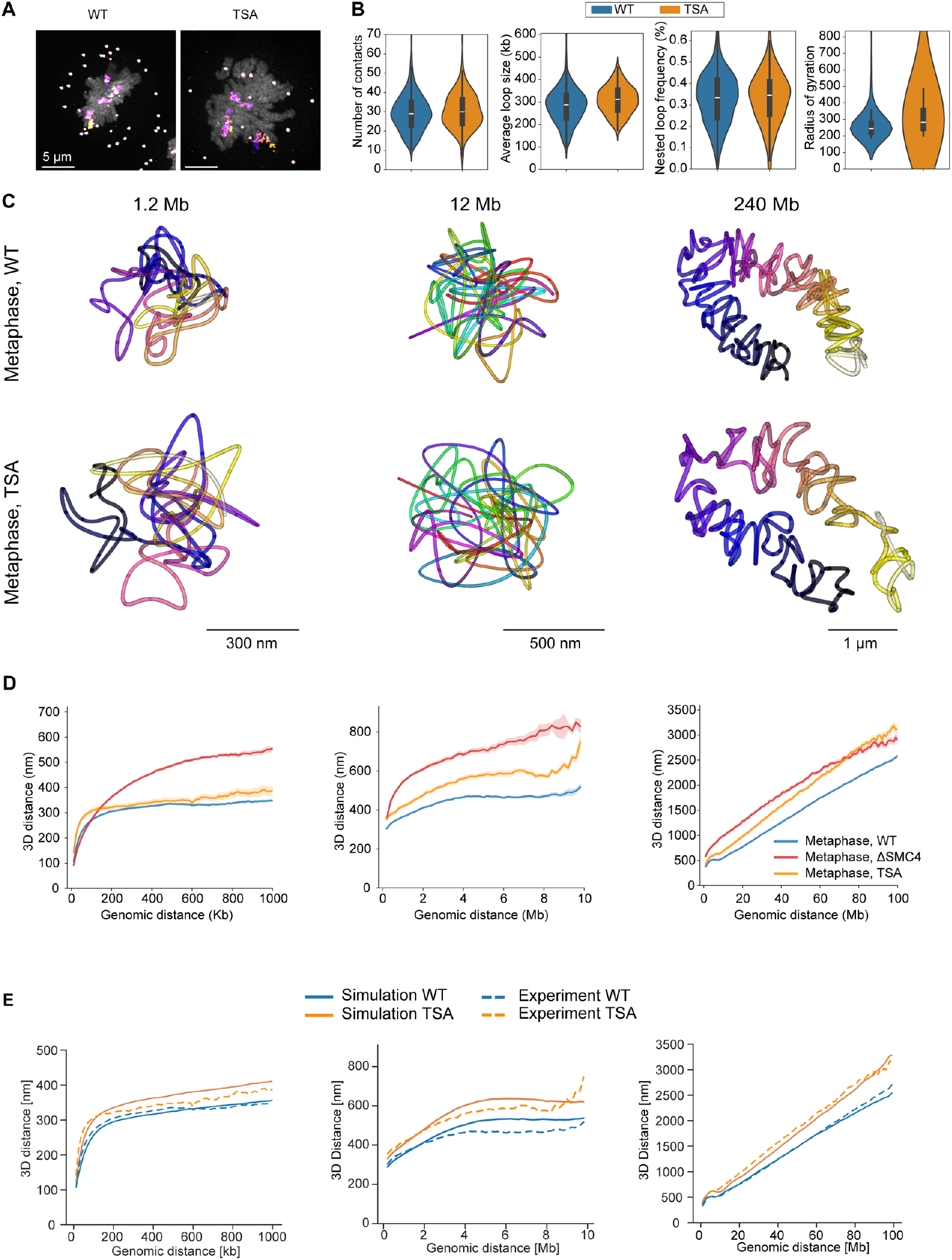
Global compaction is influenced by self-repulsion. **A** Wild type (WT) HeLa Kyto cells and HeLa Kyoto cells treated with TSA before mitotic entry. Exemplary maximum-intensity projected micrograph of metaphase chromatin (DAPI, grey) with full chromosome 2 traces (DNA-FISH, multi-coloured). Data representative of 212 WT metaphase cells from 3 independent experiments and 205 prometa-/metaphase TSA-treated cells from 2 independent experiments. **B** Trace metrics from chr5:149500723-150699962 (1.2 Mb, 12 kb resolution) for WT and TSA-treated cells. Data from 152 (398) WT cells (traces), 3 independent experiments, and 67 (168) TSA-treated cells (traces), 2 independent experiments. **C** Reconstructed traces for chromosome 5, 1 Mb scale (12 kb resolution), chromosome 2, 10 Mb scale (200 kb resolution) and whole chromosome 2 (1 Mb resolution) from WT and TSA-treated cells at metaphase. Data representative of Chr5, 1 Mb: 152 (398) WT cells (traces), 3 independent experiments, and 67 (168) TSA-treated cells, 2 independent experiment; Chr 2, 10 Mb: 159 (286) WT cells (traces), 2 independent experiments and 69 (126) TSA-treated cells (traces), 2 independent experiments; Chr2 whole: 130 (212) WT cells (traces), 3 independent experiments and 19 (31) TSA-treated cells (traces), two independent experiments. **D** Distance scaling plots for chromosome 5, 1 Mb scale, chromosome 2, 10 Mb scale and whole chromosome 2 at metaphase of TSA-treated cells together with corresponding WT and ΔSMC4 data. Data from Chr5, 1 Mb: 71 (198) TSA-treated cells (traces), 2 independent experiments; Chr2, 10 Mb: 73 (135) TSA-treated cells (traces), 2 independent experiments; Chr2, whole: 24 (69) TSA-treated cells (traces), 2 independent experiments. See Fig. 4 for details on WT and ΔSMC4 data. **E** Distance scaling plots from simulated 100 Mb WT and TSA-treated chromosomes, sampled as the corresponding experimental data (see **D** for details). TSA-treatment was simulated by increasing the repulsive potential between monomers. Simulated data from 20 dynamically simulated chromosomes per condition, sampled at metaphase (40 minutes).

## Discussion

Here, we present DNA tracing data of native mitotic chromosomes. Combining multiscale 3D DNA tracing in single cells with mechanistic exploration of data-constrained polymer simulations, we have found that the fine-scale structure and the large-scale emergent properties of mitotic chromosomes, including their characteristic rod-like shape can be explained by a simple self-organization mechanism driven by the counteracting forces of Condensin-driven loop extrusion and self-repulsion of DNA.

Extrusion of on average 6-8 Mb long Condensin II loops, that overlap with each other multiple times in a gapless manner along the whole chromosomal DNA molecule is a key structuring element of this model. This shortens the axial length of the polymer and enforces stacking of DNA when reaching the self-repulsion limit. Importantly, the progressive formation of long, overlapping loops require processive loop extrusion as long as Condensin is bound to DNA, which has been shown to occur in vitro^8,36^. Passing of Condensin motors^32^ is also key to generate gapless overlaps between loops, especially in the light of the reported unidirectional extrusion of Condensins^8^. Combined, the processivity while bound and passing allow Condensin loops to reach the average 6-8 Mb size that recapitulates the observed distance scaling minimum. The redundancy in loop overlaps also explains the robustness of mitotic chromosome shape, despite varying Condensin abundances observed between single mitotic cells^9^. Furthermore, redundantly overlapping loops provide an explanation for the mechanical resilience of mitotic chromosomes without requiring a contiguous Condensin-based axis^3,4^.

Our results are consistent with earlier contact-frequency analyses of cell populations by HiC^10,11^, and provide a fully self-consistent model for how the action of the two Condensin isoforms explain major features of both nanoscale and macroscale organization of mitotic chromatin without additional assumptions of a central axis or external confinement of chromatids^11,12^. Our model predicts that through the generation of increasingly long loops from prophase to metaphase, Condensin II effectively decreases the length of chromatids while increasing their width. In contrast, the more abundant and dynamically bound Condensin I complexes^9^ lead to smaller, short-lived loops nested inside these large loops, that induce a lateral compaction and increase the self-repulsion of the polymer. This self-repulsion in turn will constrain the maximal longitudinal compaction Condensin II can achieve. Our model thus provides a mechanistic explanation for previous observations of global changes to chromosome shape resulting from isoform-specific Condensin depletion^5,6,37^.

Previously, a higher-order regular folding of mitotic chromosomes into a helix or a perverted helix has been proposed^11–13,38–40^. While our direct 3D tracing data does not provide support for a regular helix, our polymer simulation can be used to explore under what conditions it could emerge. Key would be that Condensin II motors would extrude loops of highly reproducible length, which would result in regular undulations in pairwise distance scaling from an initial minimum (see Fig. S4D). However, this does not match the experimental data we obtained in cultured human cells. Rather, our direct 3D tracing suggests a wide distribution of Condensin II loop lengths, resulting in the characteristic distance scaling dip at the average loop size of 6-8 Mb, without the formation of a regular, handed helix in individual chromosomes (see Fig. S2E).

In conclusion, by providing the first direct visualization of the internal fold of mitotic chromosomes, we have been able to propose a realistic model of the self-organization principles of DNA that allow faithful genome inheritance. This model is constrained by quantitative experimental data and free of assumptions regarding chromosome structure. The counteracting forces of gathering the linear DNA molecule into long, overlapping loops and limiting their packing by self-repulsion are sufficient to explain the experimentally determined structures by polymer self-organization.

## Acknowledgements

We thank Prof. Narumiya, Kyoto University for sharing HeLa Kyoto cells. We thank Dr. Gerlich, IMBA, Vienna for sharing HeLa Kyoto SMC4-mAID-Halo, AAVS1-OsTir1 F74G-SNAP cells and for advice on TSA treatment protocols. We thank the ALMF, HPC and electronics workshop teams at EMBL for their generous support. We thank other members of the Ellenberg lab for fruitful discussions and advice.

## Author contributions

K.S.B. contributed to conceptualization, investigation, methodology, software, formal analysis, visualization, writing and funding acquisition. A.B. contributed to conceptualization, investigation, methodology, formal analysis, visualization, writing and funding acquisition. N.M. contributed to methodology, investigation, validation and reviewing. R.J. contributed to conceptualization, funding acquisition and reviewing. J.E. contributed to conceptualization, supervision, project administration, writing and funding acquisition. **Funding:** This work was supported by grants from the National Institutes of Health Common Fund 4D Nucleome Program (Grant U01 EB021223 / U01 DA047728) to J.E., as well as by the The Paul G. Allen Frontiers Group through the Allen Distinguished Investigator Program to J.E. and R.J.. A.B. has received a PhD fellowship from the Boehringer Ingelheim Fonds and K.S.B. was supported by the Alexander von Humboldt foundation. Funded by the European Union. Views and opinions expressed are however those of the author(s) only and do not necessarily reflect those of the European Union or the European Research Council Executive Agency. Neither the European Union nor the granting authority can be held responsible for them. This work was supported by ERC grant (MITOFOLD, 101142430). This work was supported by DFG Priority Programme “Spatial Genome Architecture in Development and Disease” (SPP 2202).

## Materials and methods

### Cell culture

HeLa Kyoto cells (HK WT: S. Narumiya (Kyoto University, Kyoto, Japan), RRID: CVCL_1922; HK SMC4-mAID-Halo, AAVS1-OsTir1 F74G-SNAP, Schneider et al., 2022) were grown in cell culture dishes (Falcon) in high-glucose DMEM (41965-062, Thermo Fisher Scientific) containing 10% FCS (10270-106, Lot. 42F2388K, Thermo Fisher Scientific), 100 U/ml penicillin-streptomycin (15140-122, Thermo Fisher Scientific) and 1 mM sodium pyruvate (11360-039, Thermo Fisher Scientific). Cells were cultured in a humidified incubator at 37°C, 5% CO_2_ unless otherwise stated. Cells were grown to a confluency of 70-90% and passaged every 2-3 days via trypsinization with 0.05% Trypsin-EDTA (25300-054, Thermo Fisher Scientific). Cell lines were regularly checked for Mycoplasma contamination and confirmed negative. To enable non-denaturing FISH via single strand-resection, cells were incubated with 40 µM BrdU/BrdC (ratio 3:1, BrdU: B5002, Sigma-Aldrich, BrdC: sc-284555, Santa Cruz Biotech) for 18-24 hours prior to fixation (see cell synchronization).

### Mitotic shake-off

For the enrichment of mitotic cells (e.g. prometa-/metaphase) in a microscopy slide, a mitotic shake-off procedure was used. Asynchronous HK cells were seeded into T-175 flasks (175 cm^2^ surface area) and grown for 18-24 hours in the presence of 40 µM BrdU/BrdC to achieve a final confluency of 70-90%. Prior to mitotic shake-off, the medium was aspirated and replaced with 12 mL fresh DMEM. Mitotic cells were detached from the cell culture dish by knocking the culture dish 5x vertically onto a table covered with 5 paper tissues. Cells were collected in a 15 mL Falcon tube and centrifuged at 90xg for 3 minutes. Cells were resuspended in 100-150 µL fresh medium to a concentration of 2-3.5 ×10^6^ cells/mL. 35 µL of enriched mitotic cells were seeded into each channel of an Ibidi glass-bottom µ-slide (80607, Ibidi), pre-coated with poly-L-lysine for better attachment of mitotic cells. After 2 minutes incubation at 37°C, cells were washed once with 1X PBS and fixed using 2.4% PFA (15710, EMS) in PBS for 15 minutes at room temperature (RT). PFA was quenched using 100 mM NH_4_Cl in PBS for 10 minutes.

### Mitotic cell synchronization in prophase or in combination with targeted perturbation

For enrichment of cells in prophase and for pre-mitotic degradation of Condensins or inhibition of histone deacetylases with trichostatin A (TSA, Sigma-Aldrich, T8552), cells were pre-synchronized using mitotic shake-off and synchronized in the next mitosis using the Cdk1 inhibitor RO-3306 (Sigma-Aldrich, SML0569).

One day prior to mitotic shake-off, cells were seeded into a T-175 flask in the absence of BrdU/BrdC to reach a confluency of 70-90%. Mitotic shake-off was performed as described above, resuspended cells were diluted to 0.6-0.9 ×10^6^ cells/mL and 35 µL were seeded into a channel of an Ibidi µ-Slide. Cells were adhered for 20-30 minutes 37°C. Subsequently, medium was exchanged to 120 µL DMEM supplemented with 40 µM BrdU/BrdC and cells were incubated for 15 hours at 37°C, 5% CO_2_. Cells were synchronized at the G2/M transition using 10 µM RO-3306 for 7-8 hours in the presence of BrdU/BrdC. During this time, perturbations were performed. To degrade Condensins SMC4-mAID-Halo cells (Schneider et.al., 2022) were treated for 3 h with 1 µM 5-Ph-IAA (30-003, BioAcademia). Inhibition of histone deacetylases was performed in HK WT cells using 5 µM TSA for 5 h. Cells were released into mitosis by washing out RO-3306 3x with fresh DMEM. Unperturbed HK WT cells were released for 12 minutes until fixation using PFA to enrich cells in prophase. ΔSMC4 cells were allowed to enter mitosis for 30 minutes in the presence of 1 µM 5-Ph-IAA until fixation to reach a metaphase-like state. TSA treated cells were allowed to enter mitosis for 30-40 minutes in the presence of 5 µM TSA until fixation. TSA-treated cells, however, failed to align in a metaphase plate as described recently^25^. Cells were permeabilized for 20 minutes in 0.25% Triton X-100 in 1X PBS at RT (T8787, Sigma-Aldrich) for immunofluorescence and non-denaturing FISH. As fiducials for image registration, samples were incubated with 0.1 µm Tetraspec beads (T7279, Thermo Fisher) diluted 1:100 in 1X PBS for 10 minutes at RT.

### Library design and amplification

oligoFISH probes were selected and filtered using oligoMiner^41^. Genome complementary sequences were 36-42 nt long with a melting temperature of 42-46°C (adjusted for 50% formamide, 2XSSC), and specificity filtered by oligoMiner linear discriminant analysis 42°C model with a probability threshold of 0.9, followed by a jellyfish 18 bp k-mer filter with k=10 (k=20 for 1 Mb regions). 20 bp sequences for PCR-amplification were appended to all probes and used during FISH to identify each full chromosome, 10 Mb or 1 Mb region. For probes in 1 Mb and 10 Mb regions, duplicate 12-bp spot-specific barcodes were appended for probes targeting subsequent 12 kb (densely tiled) or 30 kb (spaced by 200 kb) windows, while for full chromosomes each probe had a single 15 Mb segment-specific barcode and duplicate spot-specific barcode targeting each 100 kb (chromosome 2) or 30 kb (chromosome 14) region spaced by 1 Mb. For 1 Mb regions, all filtered probes were used, giving an average of 150 probes per 12 kb spot. Probes for 10 Mb regions and full chromosomes were randomly sampled so that each position had at most 200 and 170 probes respectively. All probe sequences are listed in Supplementary Table 2. Libraries were ordered as oligo pools (Genscript). 12-bp barcode sequences were designed as described before^17^. Imager probes targeting 12-bp barcodes were ordered with a 5’-azide (Metabion GmbH) and labelled with Cy3B-alkyne (AAT Bioquest) or Atto643-alkyne (Attotec) using click chemistry (ClickTech Oligo Link Kit, Baseclick GmbH) according to the manufacturer’s instructions. Oligo pools were amplified by reverse transcription and in vitro transcription as described before^17,42^.

### Immunofluorescence

Immunofluorescence of Condensin’s SMC2 subunit was performed to visualize chromatid axes relative to the DAPI/tracing signal. After permeabilization and addition of fiducial beads, cells were incubated with blocking buffer (2% BSA in 0.05% Triton X-100 in 1X PBS) for 30 minutes at RT. Following blocking, cells were incubated with the primary antibody against SMC2 (ab10412, Abcam, 1:1000) in blocking buffer overnight at 4°C or for two hours at RT. Subsequently, cells were washed 3x for 5 minutes in blocking buffer and then incubated with the AF488-labelled secondary antibody (A-11034, Molecular Probes, 1:1000) for 1h at RT in blocking buffer. Cells were washed 3x for 5 minutes in 1X PBS.

### Non-denaturing FISH

Non-denaturing FISH (RASER-FISH^21^) was performed as described recently^17,22^. Permeabilized cells with fiducial beads (and optionally immunofluorescence staining) were incubated for 15 minutes with 0.5 ng/µL DAPI (D9542, Sigma-Aldrich) and subsequently washed 2x with 1X PBS to sensitize BrdU/C-labeled cells for UV treatment. UV treatment of cells was performed by exposing the Ibidi µ-slide without a lid to 254 nm UV light for 15 min (Stratalinker 2400 fitted with 15W 254 nm bulbs, part no. G15T8) to induce single strand nicks. Nicked DNA was digested using Exonuclease III (M0206, NEB) at a final concentration of 1 U/µL in NEB buffer 1 at 37°C for 15 minutes in a humidified chamber. Cells were washed 3x with 1X PBS. Cells were post-fixed with 5 mM Bis(NHS)PEG5 (803537, Sigma-Aldrich) in 1X PBS for 30 minutes at RT to preserve sample fixation during primary probe hybridization. The sample was equilibrated with 1X hybridization buffer (50% formamide (FA, AM9342, Thermo Fisher), 10% (w/v) dextran sulfate (D8906, Sigma-Aldrich) in 2xSSC (AM9763, Thermo Fisher) for 15 minutes at 37°. Primary probe libraries were diluted to a concentration of 100-300 ng/µL in 1X hybridization buffer and hybridized for 1-2 days at 37°C for 15 minutes in a humidified chamber. After hybridization, cells were washed 2x with 50% Formamide in 2X SSC for 5 minutes at RT, 3x with 2X SSC containing 0.2% Tween and 2x with 2X SSC. Cells were incubated with 0.05 U/µL RNAse H (M0297S, NEB) for 20 min at 37 °C in RNAse H buffer (NEB) to remove RNA-DNA hybrids. After washing 3x with 2X SSC, DNA bridges were hybridized to primary FISH probes to enable the common fluorescent readout of each library targeting a specific genomic locus or chromosome. To this end, DNA bridges were diluted to 100 nM in secondary hybridization buffer (20% Ethylene Carbonate (EC, E26258, Sigma-Aldrich), 2xSSC) and allowed to hybridize to primary probes in the sample for 20 minutes at RT rocking. Subsequently, cells were washed 3x with 30% Formamide in 2X SSC for 5 minutes each at RT rocking, and 2x with 2XSSC.

### Chromatin tracing

Chromatin tracing using LoopTrace was performed as described recently^17,22^. FISH-treated cells were mounted on a Nikon TI-E2 microscope with Omicron lasers, a 100X 1.35 NA silicon oil immersion objective, a CSU-W1 SoRa spinning disk unit, a pentaband filter and an Orca Fusion CMOS camera, operated using NIS Elements 5.2.02 (Nikon). The Ibidi µ-Slide was connected to a custom-built automated fluidics robot based on a GRBL controlled CNC stage as described in^17^ (see https://git.embl.de/grp-ellenberg/tracebot) to enable automated liquid handling for sequential imaging. Prior to sequential imaging, 25-35 3D stacks of DAPI-stained nuclei (405 nm excitation), Condensins (488 nm excitation, see Immunofluorescence section) and fiducial beads (561 or 640 excitation) were acquired as reference images for cell classification. Images had a pixel size of 130 nm in xy and 300 nm in z and a total size of 149.76×149.76 µm in xy and covering a z-range of ∼20 µm. After that, fluorescent imager probes targeting either a single genomic spot within each FISH library or an entire library via a bridge probe were sequentially hybridized through the automated liquid handling system and imaged. We performed dual-colour tracing with Cy3B- and Atto643-labeled 12 bp imager probes. Imager probes were stored in a 96 well plate on the CNC stage, in addition to the buffers for washing (10 % FA, 2X SSC), stripping (30% FA, 2XSSC) and imaging (56 mg/mL glucose oxidase (G7141, Sigma-Aldrich), 3.4 mg/mL catalase (C30, Sigma-Aldrich), 1.5 mM Trolox (238813, Sigma-Aldrich), 10% Glucose, 50 mM Tris, 2X SSC pH 8.0), stored in extra 3-well plates, covered in parafilm and mounted on the CNC stage. Imaging buffer was covered with a layer of light mineral oil (330779 Sigma-Aldrich) to prevent buffer degradation over the course of a 20-hour imaging run. Imager strands and buffers were drawn with a syringe needle mounted in place of the CNC drill head. This needle was connected to the sample and a CPP1 peristaltic micropump (Jobst Technologies, Freiburg, Germany, flow rate of 1 mL/min at maximal speed) using 1 mm i.d. PTFE and silicone tubing (Bola, VWR), allowing to pull liquids out of the well plates and through the sample channel in an automated manner. Imager strands were sequentially hybridized for ∼ 2 minutes at 20 nM in 5% EC 2X SSC, followed by 1 minute washing with wash buffer. After addition of imaging buffer, the same cells as imaged in the reference stacks were imaged with same imaging dimensions in xyz in the 561 nm or 640 nm channels (100% laser power, 100 msec exposure time, triggered acquisition mode) to image Cy3B or Atto643-labelled imagers, respectively, alongside fiducial beads. After each round of imager strand hybridization and imaging, the imaged imager strands were stripped off using stripping buffer for 2 minutes and cells were washed again for 1 minute using wash buffer.

### 1 Mb and 10 Mb scale trace data analysis

Processing of acquired 1 Mb and 10 Mb-scale tracing data was performed as described in^17^ with code available at https://git.embl.de/grp-ellenberg/looptrace. In brief, nd2 image files were converted to OME-ZARR format. Sequential images were registered by cross-correlation, and residual sub-pixel drift was corrected by calculating the mean offset of 3D gaussian centroids of segmented fiducial beads. Images were deconvolved using an experimental PSF extracted from registering and averaging 200 fiducial bead 3D images. Tracing regions were identified and segmented based on regional barcodes using an empirical intensity threshold. Each tracing region was background-corrected by subtraction of a blank frame, and up to 3 potential spots were identified per frame by local peak detection. Centroids of each spot were super-localized by fitting a 3D gaussian function. After filtering for poor fits, the fit closest to the median position of all high-quality unambiguous fits was retained for each frame. Chromatic aberration between Cy3B and Atto643 signals was corrected by least square fitting of the centroid of fiducial beads imaged in both channels, and traces were assigned to nuclei classified as “interphase”, “prophase”, “prometaphase” or “metaphase” based on chromosome organization of each nucleus.

### Full chromosome trace data analysis

Image conversion, registration and deconvolution were performed as for the higher resolution regions. To reduce the required number of tracing cycles, full chromosome sequential frames included the full chromosome regional barcode, 15 Mb segment barcodes and 30 spot barcodes repeating every 30 Mb (spanning 2 segments). Decoding the combined spot and segment identity identified the genomic coordinate of each spot. Full chromosome regions were segmented from regional barcodes using a random forest classifier (scikit-learn^43^) trained on 5-10 small foreground and background regions in one field of view. All potential peaks in the region were detected on background-corrected spot frames by local peak detection in scikit-image^44^ and fit by a 3D gaussian. Fits were quality controlled and corrected for chromatic aberration as above. To assign each fit to a 15 Mb segment, the intensity of each fit coordinate was measured in each potentially matching segment frame and assigned a z-score by subtracting the mean and dividing by the standard deviation of the peak intensity in all segment frames. To account for the potential presence of FISH signal from multiple chromosomes in the same region of interest, remaining ambiguous fits (i.e. more than one high-quality fit assigned to the same genomic coordinate) were assigned by spectral clustering based on spatial and genomic position into an increasing (1-5) number of clusters until most (75^th^ quantile) fits were unambiguously assigned to a chromosome trace. Remaining ambiguous fits were removed by selecting the fits with the highest z-score. Finally, traces were assigned to the cell cycle stage of their parent nuclei.

### Trace analysis

For analysis, 1 Mb and 10 Mb-scale fits were quality-controlled based on fit standard deviation and signal to background ratio, while whole chromosome fits were used directly. Median pairwise Euclidean distances were calculated for all 3D coordinates within a single trace and used to calculate distance scaling plots. A cut-off of minimum 30 points per trace was used. Median pairwise Euclidean distances were also used to compute trace metrics for the 1 Mb regions, with a minimum cut-off of 80 points. To assess potential compaction below the highest tracing resolution, we calculated the sum of sequential 3D distances in each trace (contour length). This measure indicated that no major compaction/de-compaction happened on length scales below the sampling resolution of our chromatin tracing (Fig. S3B), validating that our 12 kb resolution sampled the major fine-scale folding changes during mitosis.

Whole chromosome axes were estimated by a 20 Mb rolling mean, and chromosome radius was calculated by the distance from each point to the closest point on the axis. Axial scaling of whole prometa- and metaphase chromosomes were calculated by robust linear regression (Siegel slopes) of physical distance scaling data from 20-60 Mb genomic distances. Loop metrics for the 1 Mb scale were based on a 100 nm 3D distance cutoff to identify loop anchors spanning at least 30 kb genomic distance, and the relative position of loop anchors determined assignment to base loop, z-loop or nested loops categories. Helicity of full chromosome traces was estimated by measuring the standard deviation of radial angles of points sampled at regular genomic intervals. 1 Mb and 10 Mb-scale traces were matched to their corresponding full chromosome traces by spatial proximity of their centroids. 3D nucleus and trace visualizations were done using mayavi after applying a rolling distance filter which removed spurious fits deviating more than 3 standard deviations from the median position of 11-point windows. Fluorescence images were displayed in napari.

### Mitotic chromosome simulations

1D loop extrusion simulations were performed using a numpy/numba based framework as described previously^17^, with loop extruder abundance and residence time set to physiological values for Condensins based on in vivo measurements in HeLa cells^9,22^. Parameters unknown in vivo (Condensin extrusion rate and interaction between Condensin complexes) were empirically tested in pilot polymer simulations. The simulations were run for 2400 one second timesteps, corresponding to ∼40 minutes mitosis from early prophase to metaphase^9^. 3D polymer simulations were performed using the polychrom package (which implements OpenMM polymer simulations^45,46^. Simulations were run on Nvidia RTX 3090 GPUs on the EMBL HPC Cluster. A harmonic potential bonded neighbouring monomers and a harmonic potential with half bond length was used for loop extruder bonds. A polynomial repulsive force between non-neighbouring monomers was used to ensure self-avoidance of the polymer. No boundary conditions were used. The simulated polymers consisted of 100 000 monomers. The simulations were equilibrated from a random walk starting conformation for 100 000 simulation steps and scaled to experimental 12 kb resolution ΔSMC4 mitotic tracing data. Dynamic mitotic simulations were performed by first equilibrating the random walk starting configuration for 100 000 simulation steps, then applying and updating loop extruder bond positions sampled from the 1D simulations every 5^th^ loop extrusion step and equilibrated for 20 000 steps, giving a total of 2 500 000 simulation steps for mitotic simulations (∼15 minute GPU run time per chromosome). In simulations of TSA-treated chromosomes the maximum repulsive potential was increased 5-fold and repulsion radius increased from 1.05 bond lengths to 1.5. The resulting polymer conformations with 1 kb monomers were sampled to match the experimental sampling of 1 Mb, 200 kb or 12 kb before analysis. Regular helical chromosomes with 12 Mb pitch were mathematically generated, and normally distributed noise with a μ=0, σ=200 nm was added to x, y, and z positions of regular helices to simulate noisy helical chromosomes.

### FISH protocol comparison

Cells from a mitotic shake-off were fixed, stained with DAPI and imaged on a spinning disk confocal. After the first round of imaging, the cells were prepared for FISH using non-denaturing or denaturing (denatured for 5 min with 0.1M HCl, then at 86 °C for 3 min during primary probe hybridization) conditions, stained with DAPI and the same cells were relocated and imaged. Mitotic cells (all stages from prophase to early telophase) were cropped manually from full fields of view of pre-treatment cells, then each pre- and post-treatment single cell image was registered in 3D using elastix with either Euler (rigid) registration or affine registration. The intensity of the pre- and post-treatment images was normalized by histogram matching, and the pixel-wise Pearson’s correlation coefficient was measured between the images using the binarized before-image as a mask.

## Data and materials availability

All chromatin tracing data generated in this study is available at https://doi.org/10.6084/m9.figshare.27003022. Custom Python code to run loop extrusion simulations and prepare polychrom polymer simulations, and additional analysis and visualization code examples are available at https://doi.org/10.6084/m9.figshare.27003022. Code used to process sequential images and extract DNA traces is available at https://git.embl.de/grp-ellenberg/looptrace. Code used to control the automated fluidics and microscopy acquisition set-up is available at https://git.embl.de/grp-ellenberg/tracebot.

## Supplementary Materials

Figs. S1 to S5

Tables S1

Movie S1

