## Supplementary Figures and Tables for "Nanoscale 3D DNA tracing reveals the mechanism of self-organization of mitotic chromosomes"

### Contents:

Supplementary Figures S1-5
Supplementary Table 1

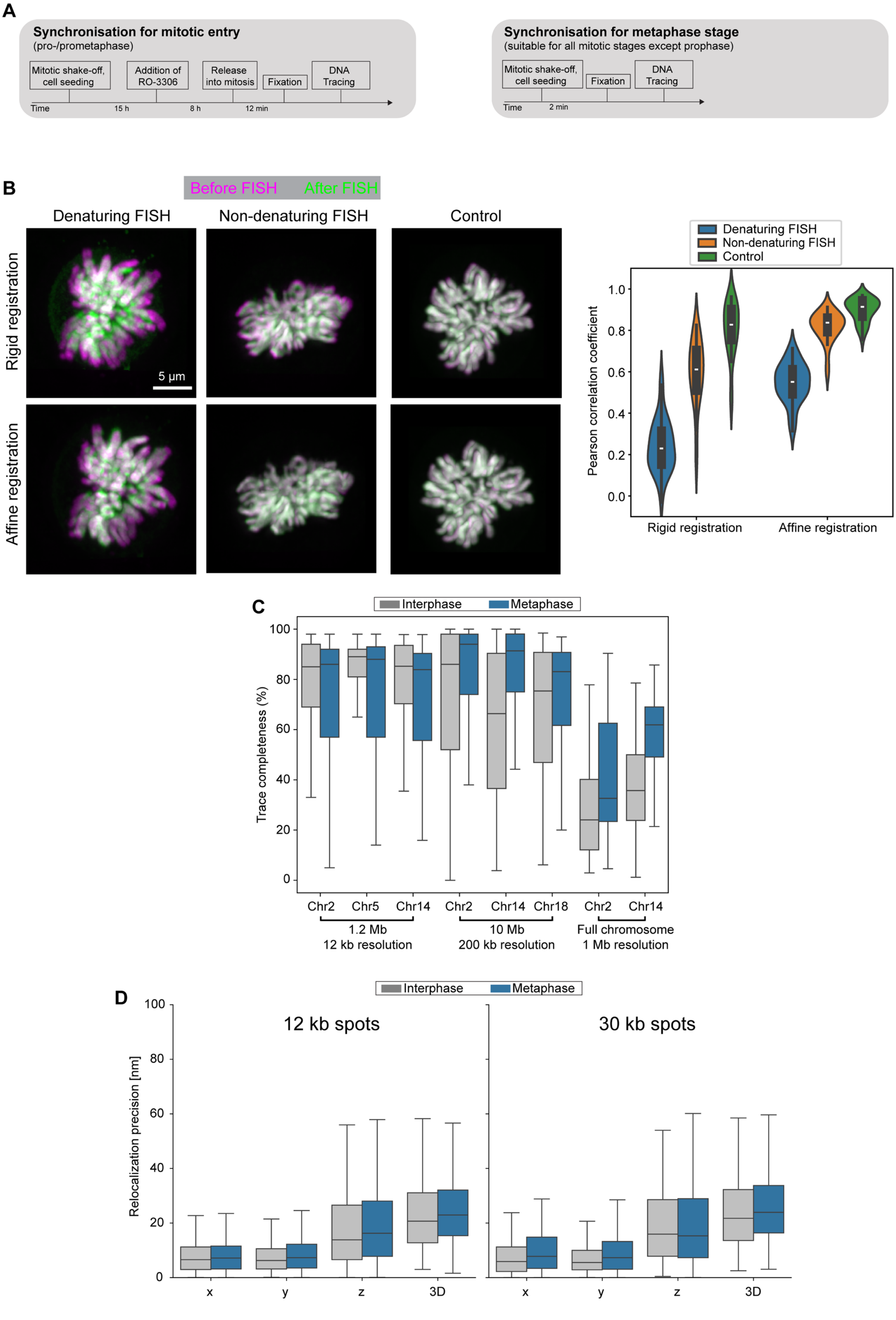
A
Synchronisation for mitotic entry
(pro-/prometaphase)
Mitotic shake-off, cell seeding
Addition of RO-3306
Release into mitosis
Fixation
DNA Tracing
Time
15 h
8 h
12 min
Synchronisation for metaphase stage
(suitable for all mitotic stages except prophase)
Mitotic shake-off, cell seeding
Fixation
DNA Tracing
Time
2 min
B
Before FISH
After FISH
Denaturing FISH
Non-denaturing FISH
Control
Rigid registration
Affine registration
5 µm
Denaturing FISH
Non-denaturing FISH
Control
Pearson correlation coefficient
Rigid registration
Affine registration
C
Interphase
Metaphase
Trace completeness (%)
Chr2
Chr5
Chr14
Chr2
Chr14
Chr18
Chr2
Chr14
1.2 Mb
12 kb resolution
10 Mb
200 kb resolution
Full chromosome
1 Mb resolution
D
Interphase
Metaphase
12 kb spots
30 kb spots
Relocalization precision [nm]
x
y
z
3D

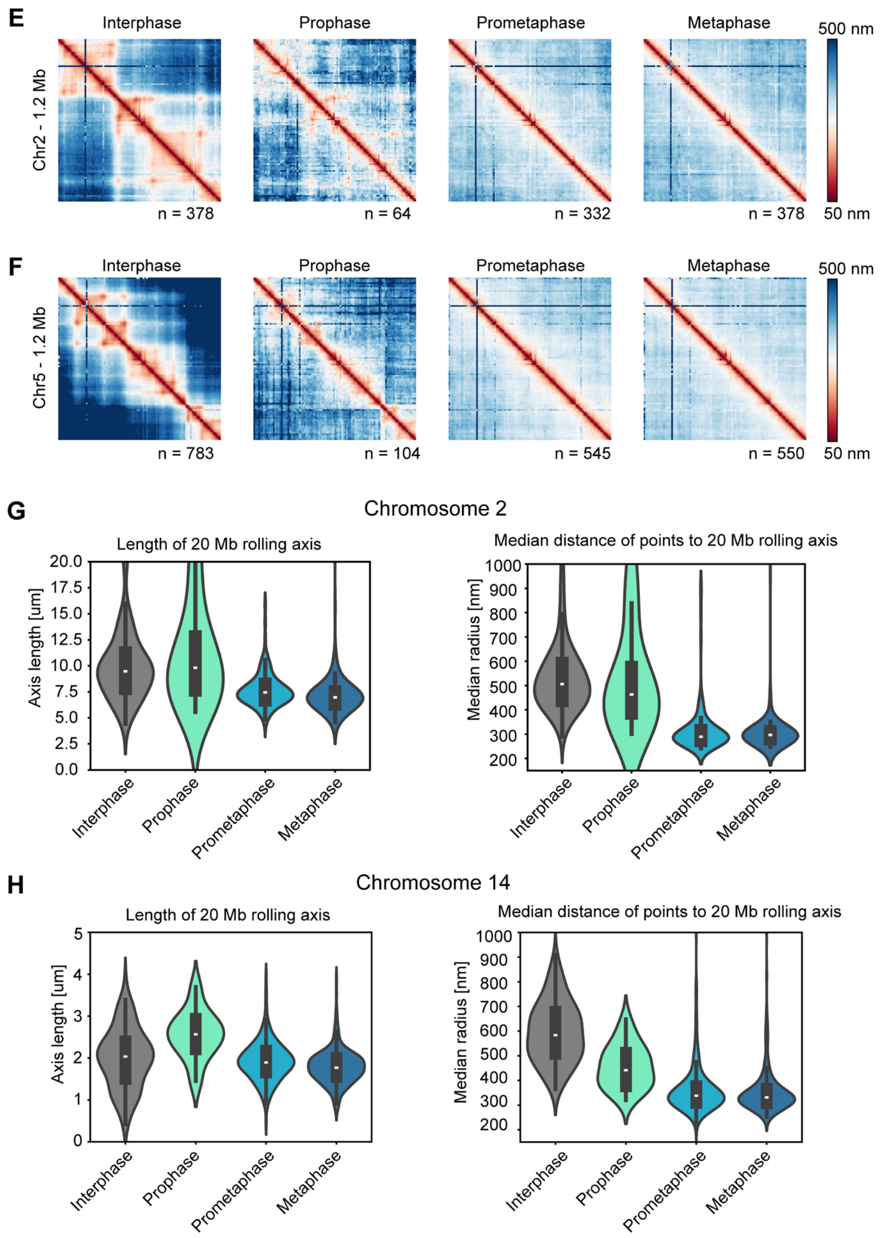


**Supplementary Figure S1**

**A** Synchronization schemes for cells enriched in pro-/prometaphase (left) and metaphase (right).

**B** Representative images (left) and pixel-wise Pearson correlation coefficient (right) of PFA-fixed mitotic cells labelled with DAPI and imaged before (magenta) and after (green) treatment with heat-denaturation FISH, non-denaturing/RASER-FISH or simulated FISH (non-denaturing FISH without library hybridization, control). Two registration algorithms were tested, with affine registration showing improved overlay accuracy as uniform scaling was better compensated. Data from one experiment with n= 92 cells (denaturing FISH), n=78 cells (non-denaturing FISH) and n=72 cells (control).

**C** Trace completeness as measured by the percentage of possible spots detected per trace across different libraries in interphase and metaphase. Data from 13455 traces in 1712 cells in 5 independent experiments. HeLa cells contain a truncated copy of chromosome 2, which reduces the completeness of whole chromosome 2 traces compared to other regions. Median, quartiles and whiskers (1.5*IQR) are shown in the plots.

**D** Tracing precision as measured by the absolute deviation in fit position after drift correction when the same genomic position was re-labelled in a different imaging cycle. Data from n=373 (interphase, 12 kb),

n=395 (metaphase, 12 kb), n=198 (interphase, 30 kb) and n=250 (metaphase, 30 kb) traces in one representative experiment of 2 independent experiments.
**E** Median pairwise distance maps of a 1.2 Mb region (Chr2:191110000-192309940) traced at 12 kb resolution. Number of traces as indicated from in total 666 cells in 2 independent experiments.
**F** Median pairwise distance maps of a 1.2 Mb region (Chr5:149500723-150699922) traced at 12 kb resolution. Number of traces as indicated from in total 874 cells in 3 independent experiments.
**G** Length and width of mitotic chromosomes, estimated by length of a 20 Mb rolling average axis and median distance of each point to the closest rolling average from chromosome 2 full chromosome library (1 Mb resolution). Data from 117 interphase, 19 prophase, 168 prometaphase and 212 metaphase chromosomes from 3 independent experiments. Median, quartiles and whiskers indicated are indicated in the plots.
**H** Data corresponding to **G** for Chromosome 14). Data from 112 interphase, 21 prophase, 292 prometaphase and 273 metaphase chromosomes from 3 independent experiments.

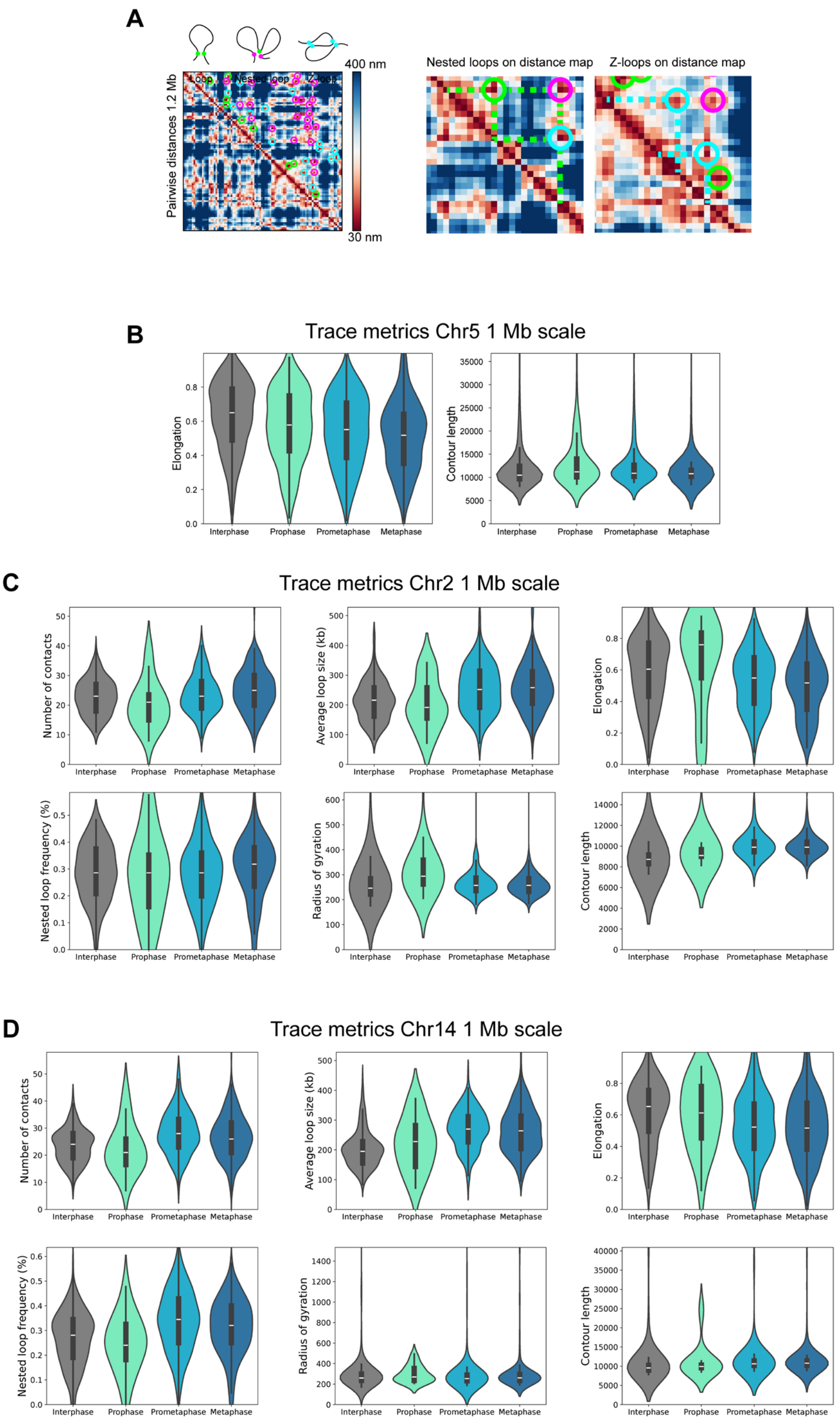
A
Loop
Nested loop
Z-loop
400 nm
30 nm
Pairwise distances 1.2 Mb
Nested loops on distance map
Z-loops on distance map
B
Trace metrics Chr5 1 Mb scale
Elongation
Contour length
Interphase
Prophase
Prometaphase
Metaphase
C
Trace metrics Chr2 1 Mb scale
Number of contacts
Average loop size (kb)
Elongation
Nested loop frequency (%)
Radius of gyration
Contour length
D
Trace metrics Chr14 1 Mb scale
Number of contacts
Average loop size (kb)
Elongation
Nested loop frequency (%)
Radius of gyration
Contour length

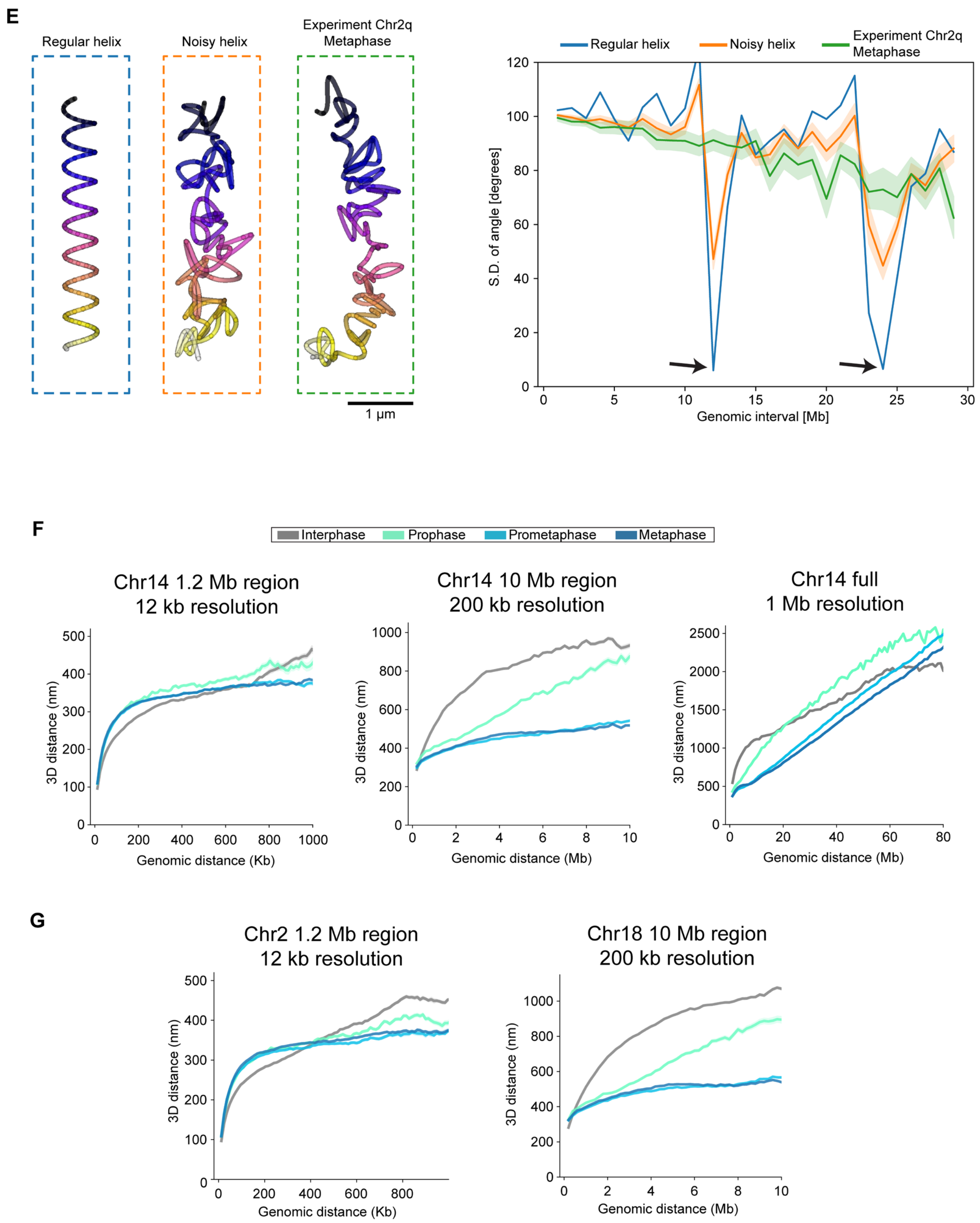


**Supplementary Figure S2**

**A** Exemplary pairwise distance map showcasing the classification of loops. Nested loops (magenta) emerge when the bases of two loops (green) merge to form a loop contact with the size of the two base loops. Z-loops (cyan) overlap with each other partially.

**B** Additional trace metrics for chr5:149500723-150699962. Elongation indicates the ratio of minor to major axis of an ellipsoid fit to the trace, while contour length measures the cumulative point-to-point length of the trace. The similar contour length for the different phases indicates that the other structural metrics that change during mitosis are well sampled at this genomic resolution. Data from

n=278 (510), n=22 (44), n=124 (297) and n=152 (398) inter-, pro-, prometa- and metaphase cells (traces) from 3 independent experiments. Median, quartiles and whiskers are shown in the plots.
**C** Trace metrics for high resolution tracing in chromosome 2:191110000-192309940. Data from n=171 (246), n=13 (19), n=100 (180) and n=134 (248) inter-, pro-, prometa- and metaphase cells (traces) from 2 independent experiments.
**D** Trace metrics for high resolution tracing in chromosome 14:50923646-52104342. Data from n=136 (189), n=9 (15), n=94 (160) and n=118 (210) inter-, pro-, prometa- and metaphase cells (traces) from one experiment.
**E** Representative examples of a simulated regular helix (left), a regular helix with added gaussian noise (center) and experimental data from chromosome 2q (right), and plotted estimation of helical regularity. Helical regularity was measured by the standard deviation in radial angle between points separated by different genomic intervals. The radial angle was measured compared to a 20 Mb rolling average axis. Local minima in the standard deviation indicating helical regularity are indicated by arrows. Data from 100 generated regular or noisy helices and the 100 best resolved chromosome 2q traces from 3 independent experiments. Mean and 95% confidence interval (estimated by bootstrapping) shown.
**F** Pairwise distance scaling plots for chr14:50923646-52104342 (12 kb resolution), chr14:45200003-56429971 (200 kb resolution) and chr14:20000036-105029664 (1 Mb resolution). Data from n=677 (1243) cells (traces), 2 independent experiments; n=645 (1201) cells (traces), 2 independent experiments and n=543 (1039) cells (traces), 2 independent experiments, respectively. Plots show median ± standard error of the mean.
**G** Pairwise distance scaling plots for chr2:191110000-192309940 (12 kb resolution), chr18: 50000077-62829816 (200 kb resolution). Data from n=711 (1280) cells (traces), 2 independent experiments and n=633 (1134) cells (traces), 2 independent experiments, respectively. Plots show median ± standard error of the mean.

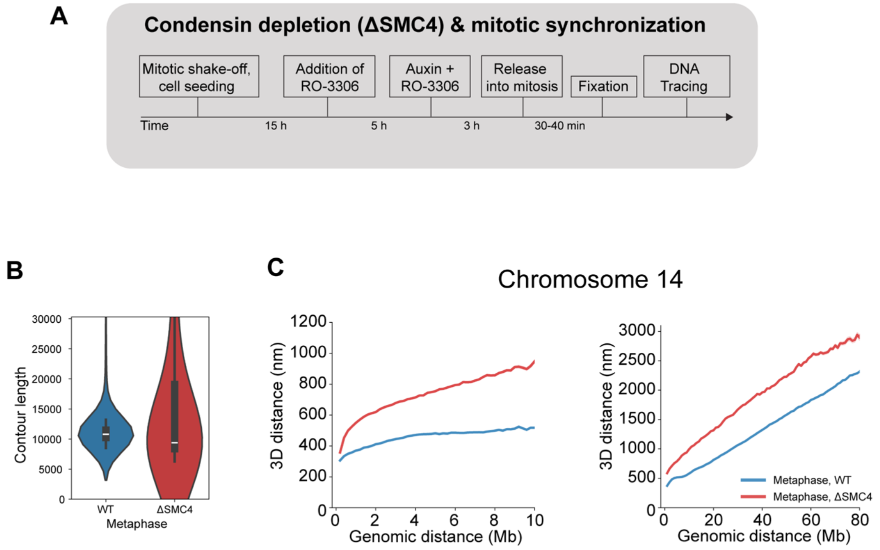


**Supplementary Figure S3**

**A** Experimental scheme for Condensin-depleted mitotic chromosomes, using the HK SMC4-mAID cell line (Schneider et al., 2022).

**B** Trace metric from chr5:149500723-150699962 (1.2 Mb, 12 kb resolution) for WT and ΔSMC4 cells. Data from 152 (398) WT cells (traces), 3 independent experiments, and 153 (265) ΔSMC4 cells, 2 independent experiments.

**C** Distance scaling plots for chromosome 14 10 Mb and whole chromosome scales. Data from Chr14, 10 Mb: 218 (434) WT cells (traces), 2 independent experiments and 217 (419) ΔSMC4 cells (traces), 2 independent experiments; Chr14, whole: 174 (357) WT cells (traces), 2 independent experiments and 78 (146) ΔSMC4 cells (traces), one experiment.

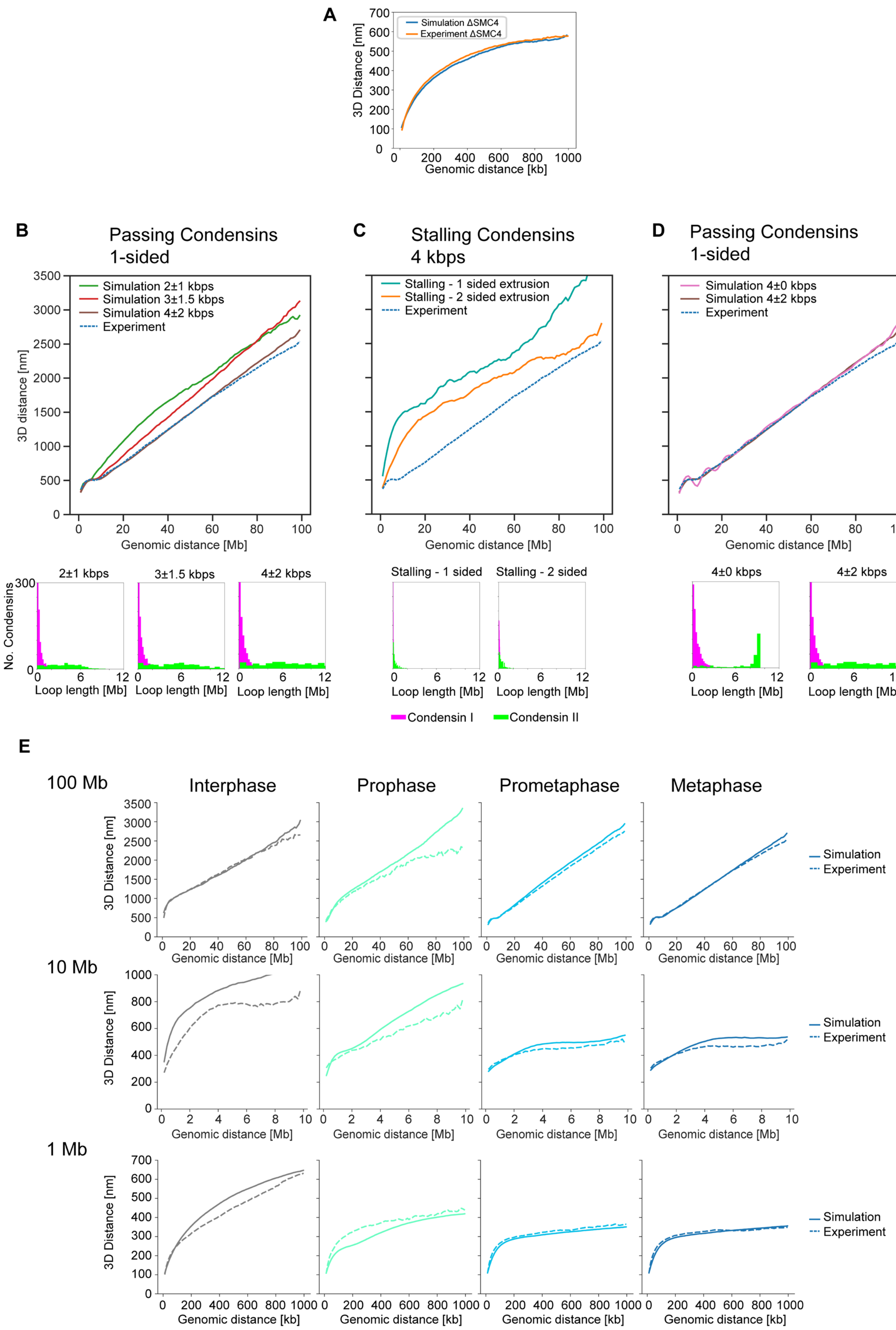
A
Simulation ΔSMC4
Experiment ΔSMC4
3D Distance [nm]
Genomic distance [kb]
B
Passing Condensins 1-sided
Simulation 2±1 kbps
Simulation 3±1.5 kbps
Simulation 4±2 kbps
Experiment
3D distance [nm]
Genomic distance [Mb]
C
Stalling Condensins 4 kbps
Stalling - 1 sided extrusion
Stalling - 2 sided extrusion
Experiment
D
Passing Condensins 1-sided
Simulation 4±0 kbps
Simulation 4±2 kbps
Experiment
2±1 kbps
3±1.5 kbps
4±2 kbps
No. Condensins
Loop length [Mb]
Stalling - 1 sided
Stalling - 2 sided
4±0 kbps
4±2 kbps
Condensin I
Condensin II
E
100 Mb
10 Mb
1 Mb
Interphase
Prophase
Prometaphase
Metaphase
3D Distance [nm]
Genomic distance [Mb]
Genomic distance [kb]
Simulation
Experiment

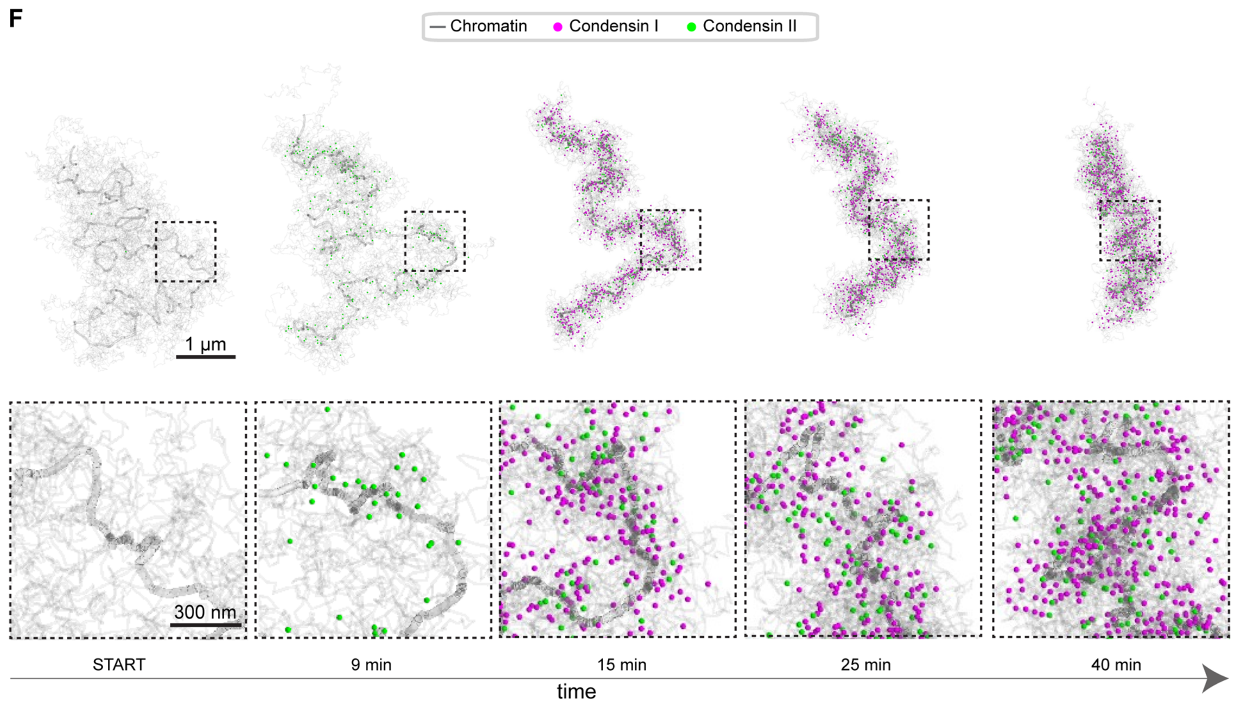


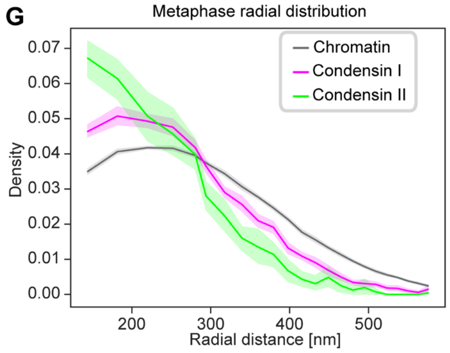


**Supplementary Figure S4**

**A** Distance scaling plot of an unconstrained polymer to experimental ΔSMC4 metaphase high-resolution tracing data (12 kb resolution, Chr5) gave 12 nm/kb monomer and 1 $k_bT$ repulsive potential as optimal polymer simulation parameters. Experimental data from 212 (608) ΔSMC4 cells (traces), 2 independent experiments.

**B-D** Distance scaling plots and histograms of Condensin I (magenta) and Condensin II (green) loop lengths resulting from alternative loop extrusion models, including (**B**) altered extrusion speed, assuming one-sided extrusion and no stalling, (**C**) One- and two-sided extrusion at 4±2 kbps with full stalling, and altered extrusion speed distributions assuming one-sided extrusion and no stalling. Simulation positions were sampled as experimental data (100 kb spots, 1 Mb resolution) ten times per 100 Mb chromosome with offset genomic positions. Simulated data from 10-20 dynamically simulated metaphase (40 minutes) chromosomes per condition. Experimental data from Chr2q, 100 Mb: n=212 (686) cell (traces), 3 independent experiments.

**E** Distance scaling plots of simulated chromosomes, assuming one-sided extrusion at 4±2 kbps and no stalling, and corresponding experimental data. Simulated traces were spatially sampled as experimental data (12 kb tiled probes, 30 kb probes with 200 kb resolution and 100 kb probes with 1 Mb resolution). Sampling timepoints were 0 min, 10 min, 32 min and 40 min for inter-, pro-, prometa- and metaphase, respectively. Simulated data from n=20 dynamically simulated 100 Mb chromosomes. Experimental data from HeLa WT cells, see Fig. 3 and Fig. S3 for detailed information.

**F** Ground-truth positions of Condensin I (magenta) and Condensin II (green) in simulated chromatin (2 kb sampling in light grey, 1 Mb rolling average in dark grey), corresponding to the example in Fig. 5A.
**G** Radial distribution of Condensin I, II and chromatin in simulated 100 Mb chromosomes. Radial distances were calculated compared to a 20 Mb rolling average axis for 20 dynamically simulated chromosomes sampled at metaphase (40 minutes), assuming one-sided extrusion at 4±2 kbps and no Condensin stalling.

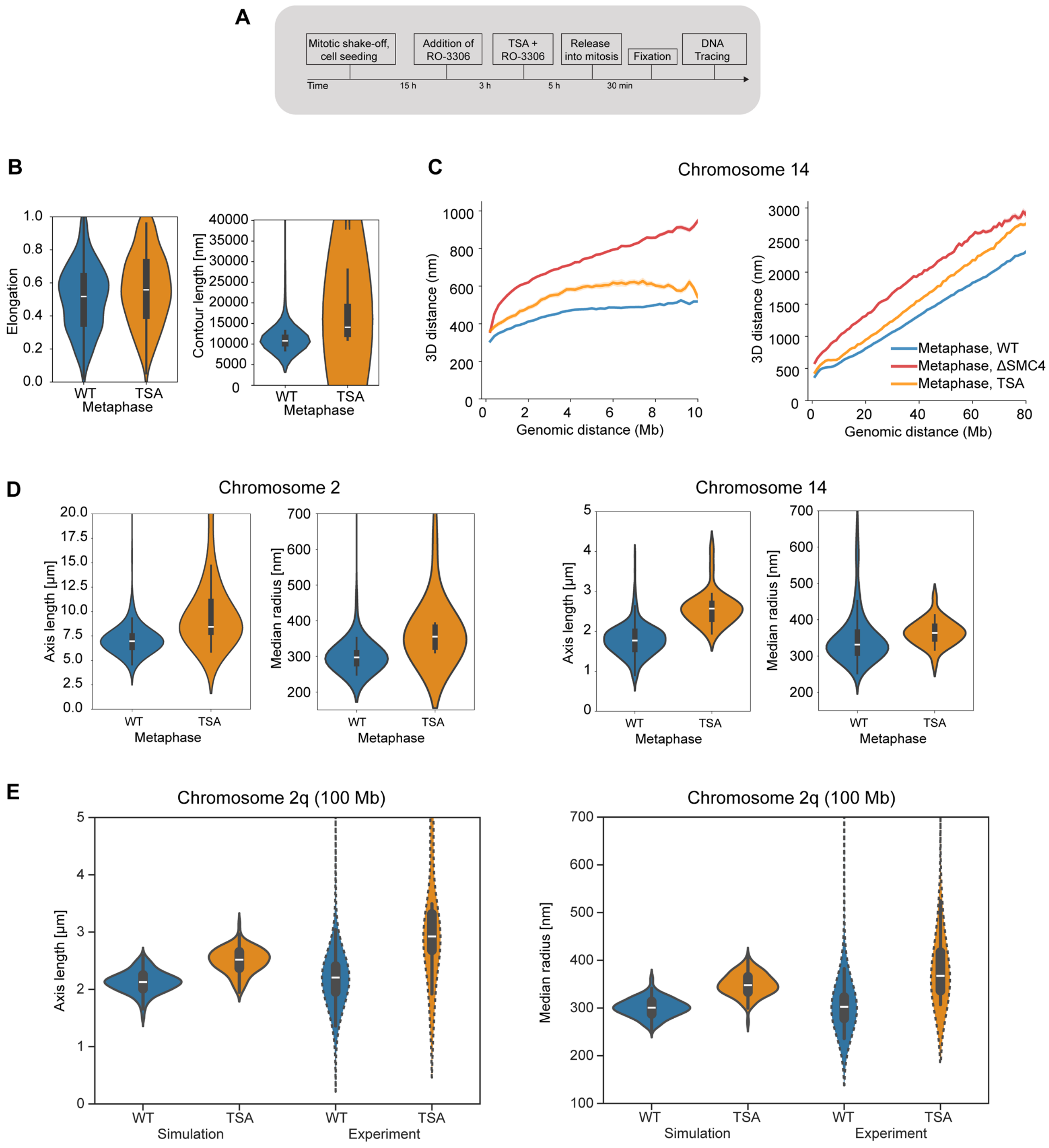


**Supplementary Figure S5**

**A** Experiment scheme for the generation of hyperacetylated mitotic chromosomes using TSA treatment during interphase.

**B** Trace metric from chr5:149500723-150699962 (1.2 Mb, 12 kb resolution) for WT and TSA-treated cells. Data from 152 (398) WT cells (traces), 3 independent experiments, and 67 (168) TSA-treated cells (traces), 2 independent experiments.

**C** Distance scaling plots from Chr 14, 10 Mb and whole chromosome scales for WT, ΔSMC4 and TSA-treated cells at metaphase. Chr14, 10 Mb: 69 (133) TSA-treated cells (traces), 2 independent experiments; Chr14, whole: 15 (30) TSA-treated cells (traces), one experiment. See Fig. S4 for details on WT and ΔSMC4 data.

**D** Length and width of WT and TSA-treated metaphase chromosomes 2 and 14, estimated by length of a 20 Mb rolling average axis and median distance of each point to the closest rolling average point from

chromosome 2 whole chromosome traces (1 Mb resolution). Data from 130 (212) WT cells (traces), 3 independent experiments and 19 (31) TSA-treated cells (traces), 2 independent experiments.
**E** Length and width of 100 Mb simulated chromosomes under WT and TSA-treated conditions, and corresponding experimental data from chromosome 2 q-arm (100 Mb). Data from 20 dynamically simulated 100 Mb chromosomes per conditions, sampled at metaphase (40 minutes). Experimental data detailed in **D**.

### Supplementary Table S1:

#### Loop extrusion simulation parameters

| Parameter | Value | Reference |
|---|---|---|
| Condensin-I abundance | 9.6/Mb | [9,22] |
| Condensin-II abundance | 2.4/Mb | [9,22] |
| Condensin-I residence time (prophase) | 0 s | [9] |
| Condensin-I residence time (prometa-/metaphase) | 150 s | [9] |
| Condensin-II residence time (pro-/prometa-/metaphase) | 3600 s | [9] |
| Extrusion rate | 1-4 ± 0-2 kbps, see main text | [32] |
| (A)symmetric extrusion | One- or two-sided, see main text | [8,32] |
| Stall probability | 0-100%, see main text | [32] |
| Duration prophase | 10 min | [9,31] |
| Duration prometa- and metaphase | 30 min | [9,31] |
| Simulation time-step | 1 s | This work |

#### Polymer simulation parameters

| Parameter | Value | Reference |
|---|---|---|
| Number of monomers | 100 000 (defined to 1kb/monomer) | This work |
| Chain/loop bond type | Harmonic bond | Polychrom documentation |
| Chain bond distance | 1 (scaled to 12 nm in analysis) | This work |
| Chain bond standard deviation | 10 % of chain bond | Polychrom documentation |
| Loop bond distance | 50 % of chain bond | Polychrom documentation |
| Loop bond standard deviation | 20 % of chain bond | Polychrom documentation |
| Periodic boundary conditions | Not used | This work |
| Collision rate | 0.03 | Polychrom documentation |
| Error tolerance | 0.01 | Polychrom documentation |
| Angle force | 1.5 | Polychrom documentation |
| Repulsive force type | Polynomial repulsive | Polychrom documentation |
| Repulsive force WT | 1 | Polychrom documentation |
| Repulsive radius WT | 1.05 | Polychrom documentation |
| Repulsive force TSA | 10 | This work |
| Repulsive radius TSA | 1.75 | This work |
| Initial polymer equilibration steps | 100 000 steps | This work |
| MD steps per loop extrusion step | 20 000 steps | This work |
| Sampling frequency of loop extrusion steps | Every 5 seconds | This work |
